# Ionizable networks mediate pH-dependent allostery in SH2 signaling proteins

**DOI:** 10.1101/2024.08.21.608875

**Authors:** Papa Kobina Van Dyck, Luke Piszkin, Elijah A. Gorski, Eduarda Tartarella Nascimento, Joshua A. Abebe, Logan M. Hoffmann, Jeffrey W. Peng, Katharine A. White

## Abstract

Intracellular pH dynamics regulate normal cell biology, but most molecular drivers remain unknown. We developed a computational pipeline to identify pH-sensitive proteins and their mechanisms. We applied the pipeline to SHP2, a pH-sensitive signaling protein, with unknown mechanism. We found that SHP2 phosphatase activity is pH-sensitive in vitro and in cells, and mutation of identified His116 and Glu252 abolishes pH-sensitive function. We also discovered that Src is an unrecognized pH-dependent kinase and mutation of the identified ionizable network abolishes pH-sensitive activity. Importantly, we found that Src kinase activity was pH sensitive even in the presence of EGF and with a phosphomimetic (Src-Y527E) mutant required for auto-inhibitory SH2 domain binding. Thus, the identified pH-sensitive regulation of Src kinase activity functions in concert with established mechanisms of Src regulation by phosphorylation. Constant pH molecular dynamics simulations performed on both SHP2 and Src support allosteric regulation mediated by pH-dependent binding of inhibitory SH2 domains to the respective catalytic domains. Finally, we show that evolutionarily conserved putative pH-sensing networks were identified across SH2 domain-containing signaling proteins. Taken together, our computational, biophysical, and cellular analyses reveal a role for pHi dynamics in allosterically regulating activity of modular SH2 signaling proteins to control biology.

**One-sentence summary:** This paper investigates the role of pH in the allosteric regulation of signaling proteins with SH2 domains, specifically focusing on SHP2 and Src

## Introduction

Transient intracellular pH (pHi) dynamics(*1*) regulate mammalian proliferation(*2*, *3*), migration(*4*), and differentiation(*5*). However, for many pH-dependent cell processes, the molecular mediators are unknown(*6*). All proteins have ionizable residues that can bind or release protons, thereby changing their charged state. For an ionizable residue to have pH-dependent biological function, the H^+^ binding affinity (pK_a_) of one or more residues must be in the physiological pH range (6.8-8.0). It is not surprising that most cytoplasmic pH-sensitive proteins (pH sensors) with characterized molecular mechanisms function through titratable histidine residues (solution pK_a_ of 6.5). However, the pK_a_s of ionizable residues are sensitive to electrostatic microenvironment(*7*). Thus, local protein microenvironment can up- or down-shift pK_a_s of glutamates, aspartates, and lysines into the physiological pH range(*8–12*). Further complicating this pH sensing landscape, ionizable residues can cooperatively regulate pH-dependent membrane proteins(*13*, *14*) and can be sequestered within cytoplasmic protein cores or buried at the interface of protein-protein complexes(*15*). However, complex networks of ionizable residues are difficult to identify *a priori* based on structure-gazing, and thus many cytoplasmic proteins known to have pH-sensitive function have unknown or incompletely characterized molecular mechanisms.

## Results

### Pipeline to identify putative pH-sensitive networks

To more quickly identify pH-sensitive proteins and characterize unknown pH sensing mechanisms, we developed an in silico prediction pipeline to identify putative pH sensing nodes based on 3D protein structure (**Fig. 1A**, see methods for details).

**Figure 1.**
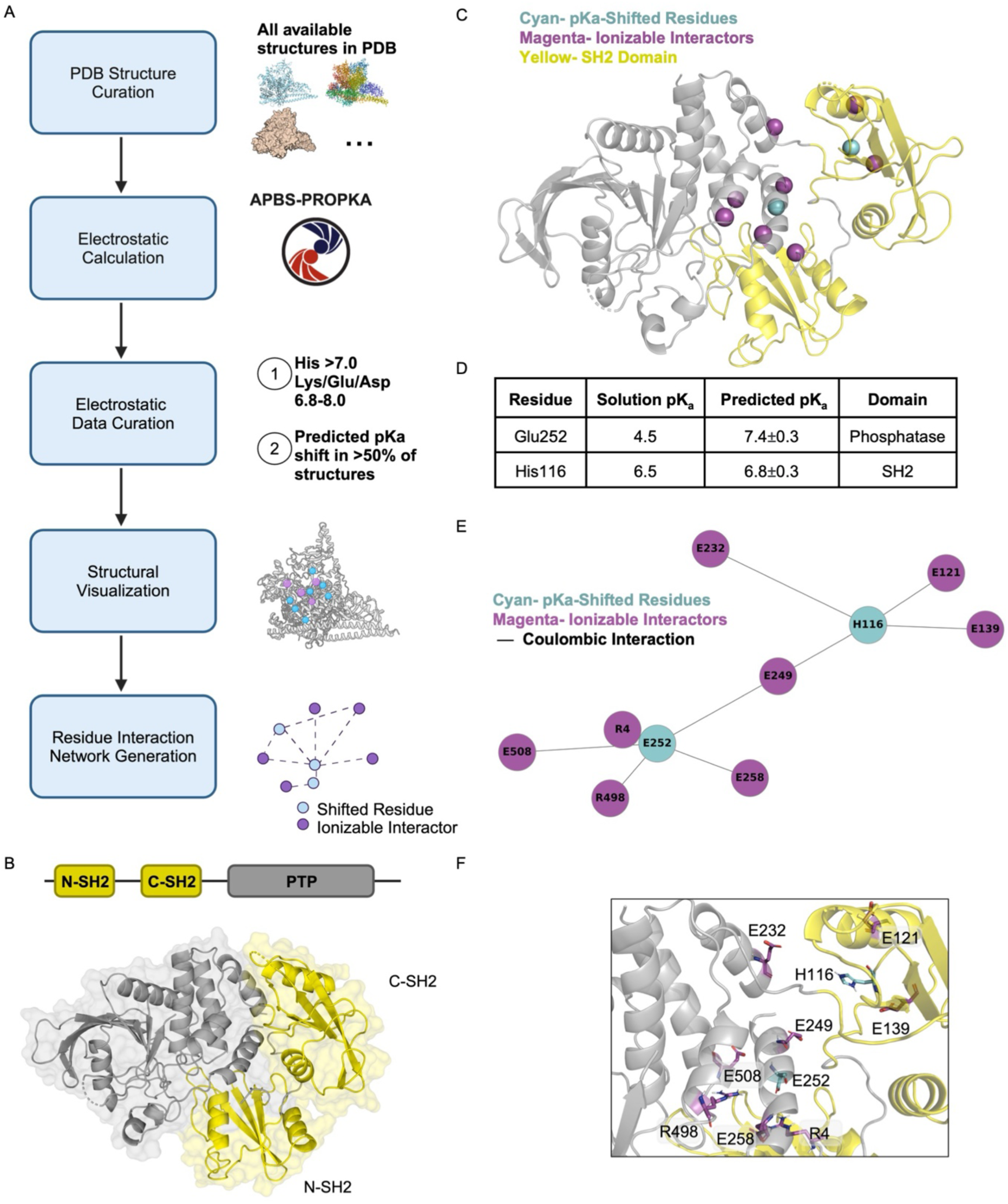
New pipeline to identify pH-sensitive networks predicts SHP2 His116 and Glu252 have physiological pK_a_s. (A) Schematic for in silico pK_a_ prediction method for proteins with solved structures. Briefly, all available structures in the protein database are curated, electrostatic properties are calculated using PROPKA, results are filtered for ionizable residues with physiologically relevant predicted pK_a_ values, and data are visualized in a 3D structure or a 2D residue interaction network. (B) Crystal structure of SHP2 shown in cartoon and surface format (PDB ID:2SHP). Protein tyrosine phosphatase (PTP) domain colored in grey, SH2 domains colored in yellow. (C) Structure of SHP2 (PDB ID:2SHP) with protein tyrosine phosphatase (PTP) domain in grey and SH2 domains in yellow. Residues identified through in silico ionizable network prediction pipeline shown in spheres. Residues with predicted pK_a_ shifts (cyan) cluster with ionizable interactors (magenta) across the phosphatase-SH2 domain interaction interface of SHP2. (D) Table of predicted pK_a_s for cyan residues identified using *in silico* ionizable network prediction pipeline on 47 SHP2 structures (mean ± SD). (E) Residue interaction network of residues with predicted pK_a_ shifts (cyan) and their ionizable interactors (magenta). Length of edges reflect the strength of the coulombic interaction, with stronger coulombic interactions having shorter edge lengths (F) Zoom of SHP2 structure at the PTP-SH2 interaction interface. Networked residues from C and E are shown in stick. Residues with predicted pK_a_ shifts in cyan and ionizable interactors in magenta.

Briefly, we use PROPKA(*16*) to predict the pK_a_s of ionizable groups based on available structure(s) of a protein of interest. Next, we filter these results for buried ionizable residues with physiological pK_a_s and generate 2D and 3D maps showing the identified residue networks and other ionizable coulombic interactors. We can perform this pipeline on multiple full-length structures of the protein of interest, increasing the likelihood of identifying functional networks. Through the generation of interaction network maps and structural maps, our pipeline provides meaningful perspectives on the functional roles of pH-sensing residues based on their location and clustering within the protein.

Our pipeline leverages structural flexibility through the analysis of multiple solved protein structures and integrates PROPKA3 for efficient pKa estimation. Previous studies collected experimental pK_a_ values from a diverse set 34 proteins with 290 experimentally determined pK_a_s for Asp, Glu, His, Tyr and Lys residues (*17*). Using our pipeline, we calculated the pK_a_s for these previously determined ionizable residues and found that our pipeline achieves high accuracy, with a correlation coefficient of 0.954 and a root-mean-square deviation (RMSD) of 0.888 (**Fig. S1A, B**). The majority of proteins in this dataset had pK_a_ shifts of less than 1.5 pH units, with just a few residues exhibiting shifts of up to 3.5 pH units (*17*). The Garcia-Moreno lab previously used NMR to measure pK_a_s of buried ionizable residues in Staphylococcal nuclease (SNase) mutants, and found large shifts in pKas (up to 5 pH units) from their solution pK_a_ values (*18–20*). To determine whether our pipeline can accurately predict large pK_a_ shifts, we compared the delta pK_a_s of the SNase NMR dataset against our predictive pipeline and found that our pipeline generally had strong correlation values with an overall RMSD of 1.6 pH units (**Fig. S1C, D**). Furthermore, we found that the pipeline accurately identified whether the residue was far upshifted or far downshifted. Importantly, there was no bias in pKa calculation errors using our pipeline for histidines, aspartates, and glutamates compared to experimental pKas in the SNase dataset. Our pipeline consistently under-predicted lysine pKa shifts compared to experimental pKas determined for the SNase dataset (**Fig. S1C**), though no such bias was found for lysine residues in the more diverse protein dataset (**Fig. S2A**).

The performance of our pipeline compares favorably to prior work developing a computationally complex pKa prediction programs (Rosetta pH, RMSD 0.85) (*17*). While multi-conformation continuum electrostatic (MCCE) calculations have been frequently used to analyze pKa shifts based on protein structures (*21*, *22*), MCCE is computationally expensive and does not achieve improved RMSDs compared to our computationally inexpensive approach. RMSDs from “quick calculation” MCCE are generally >1.3 pH units, and calculations including rotamers and side chain optimization are required to achieve RMSDs <1.0 pH unit by MCCE (*21*). Similar computationally efficient approaches have been developed to predict protein pKas such as pHinder (*13*) and BRIDGE (*23*), both arising from work on membrane proteins. Unlike these approaches, our approach is agnostic to protein localization and does not require a path for structural waters from bulk solvent to the identified buried ionizable networks(*13*, *24*).

Thus, our computational pipeline offers a comprehensive and efficient approach to identify residues with predicted physiological pKas that may function as pH-sensitive residues in proteins. Through the generation of interaction network maps and structural maps (**Fig. 1A**), our pipeline provides meaningful information to identify putative pH-sensing residues and link them to protein function based on their location and clustering within the protein. We next applied our pipeline to predict pH-sensing mechanisms and identify novel pH sensitive proteins.

### Analysis pipeline applied to SHP2 identifies His116 and Glu252 with predicted upshifted pK_a_s

We first applied our analysis pipeline to the SH2 domain-containing signaling protein SHP2. SHP2 is a regulator of multiple pH-sensitive processes including cell proliferation, migration, differentiation, growth, and survival(*25–27*). SHP2 has also been shown to have pH-dependent phosphatase activity in vitro, with optimal activity at pH 7.0 and decreasing activity both below and above 7.0(*28*), suggestive of multi-group titration. While SHP2 activity is pH sensitive in vitro, the pH-dependent molecular mechanism has not been identified. Furthermore, prior work showed that SHP2 activity positively regulates endothelial cell migration and angiogenesis and vascular smooth muscle cell function(*26*), but the role for pH-sensitive SHP2 function in cells is less clear.

SHP2 consists of three globular domains: a protein tyrosine phosphatase (PTP) domain responsible for catalyzing the dephosphorylation of tyrosine-phosphorylated proteins, and two Src Homology 2 (SH2) domains that serve as phosphotyrosine (pTyr)-recognition domains(*29*) (**Fig. 1B**). In the absence of a tyrosine-phosphorylated binding partner, the N-terminal SH2 domain binds directly to the phosphatase domain, blocking the active site(*30*). Upon ligand-induced receptor activation or signaling events, the SH2-PTP interaction is disrupted when the SH2 domains are displaced by a tyrosine phosphorylated binding partner(*30*). Thus, the SH2 domains control SHP2 activity by allosterically inhibiting the PTP domain in the bound state and influencing SHP2 localization via direct interactions with phosphorylated binding partners when signaling active(*30*).

Because SHP2 has no known molecular mechanism of pH-sensing, our goal was to identify the putative pH sensing nodes driving SHP2 pH-dependent phosphatase activity. We processed 47 structures of SHP2 available in the PDB through the computational pipeline. Importantly, we did not filter the structural input data: SHP2 structures included active conformations, inhibited conformations, and SHP2 proteins with gain- or loss-of-function mutations. Our analysis revealed a conserved network of ionizable residues clustering at the interface between the SH2 and PTP domains of SHP2 (**Fig. 1C**). This residue network contained two residues with predicted physiological pK_a_s: His116 and Glu252, with predicted pK_a_s of 6.8±0.3 and 7.4±0.3, respectively (**Fig. 1D**). His116 is in the C-SH2 domain and directly interacts with a cluster of glutamate residues (E121, E139, E232, and E249). Similar glutamate “cages” have been identified in other wild-type pH sensors including talin(*31*) and RasGRP1(*32*). Glu252 is in the phosphatase domain at the interaction interface with the N-SH2 domain and interacts with R4, E249, E258, R498, and E508 (**Fig. 1E, F**). We next determined the structural conservation of the identified residues using Consurf(*33*). Analyses of structural conservation can frequently reveal functionally important regions, such as catalytic and ligand-binding sites(*33*). We found that the C-SH2 and N-SH2 are both highly conserved at the interaction interface with the PTP domain (**Fig. S2A**). We also determined that His116 and Glu252 are sequence-conserved across SHP2 from lower-order organisms (**Fig. S2A-C**). Taken together, these data suggest that the SH2 domain binding interface in SHP2 has an evolutionarily conserved network of ionizable residues with predicted physiological pK_a_s.

### SHP2 pH sensitive function requires both His116 and Glu252

We hypothesized that the previously uncharacterized pH-dependent molecular mechanism of SHP2 requires the ionizable network identified using our computational pipeline. We first assessed the in vitro phosphatase activity of wild-type SHP2 under different buffer pH (pH 6.1-8.0) using a generic para-nitrophenyl phosphate (PNPP) substrate. We found that wild-type SHP2 phosphatase activity is pH dependent, with maximal activity at pH 6.7 and reduced activity both above and below this value (**Fig. 2A, C, S3A**). We found that k*_cat_* increased from pH 6.1 through 6.7 and decreased from 7.0 through 8.0 forming a biphasic activity curve **(Fig. 2C)**, while K_m_ for the PNPP was pH-insensitive (**Fig. S3A**). This result confirmed previous in vitro characterization of SHP2 activity(*28*), showing similar apparent pK_a_s of ∼6.7 and ∼7.1 in both the k*_cat_* (**Fig. 2A, C, S3B**) and catalytic efficiency (k*_cat_*/K_m_) pH profiles (**Fig. S3C**).

**Figure 2.**
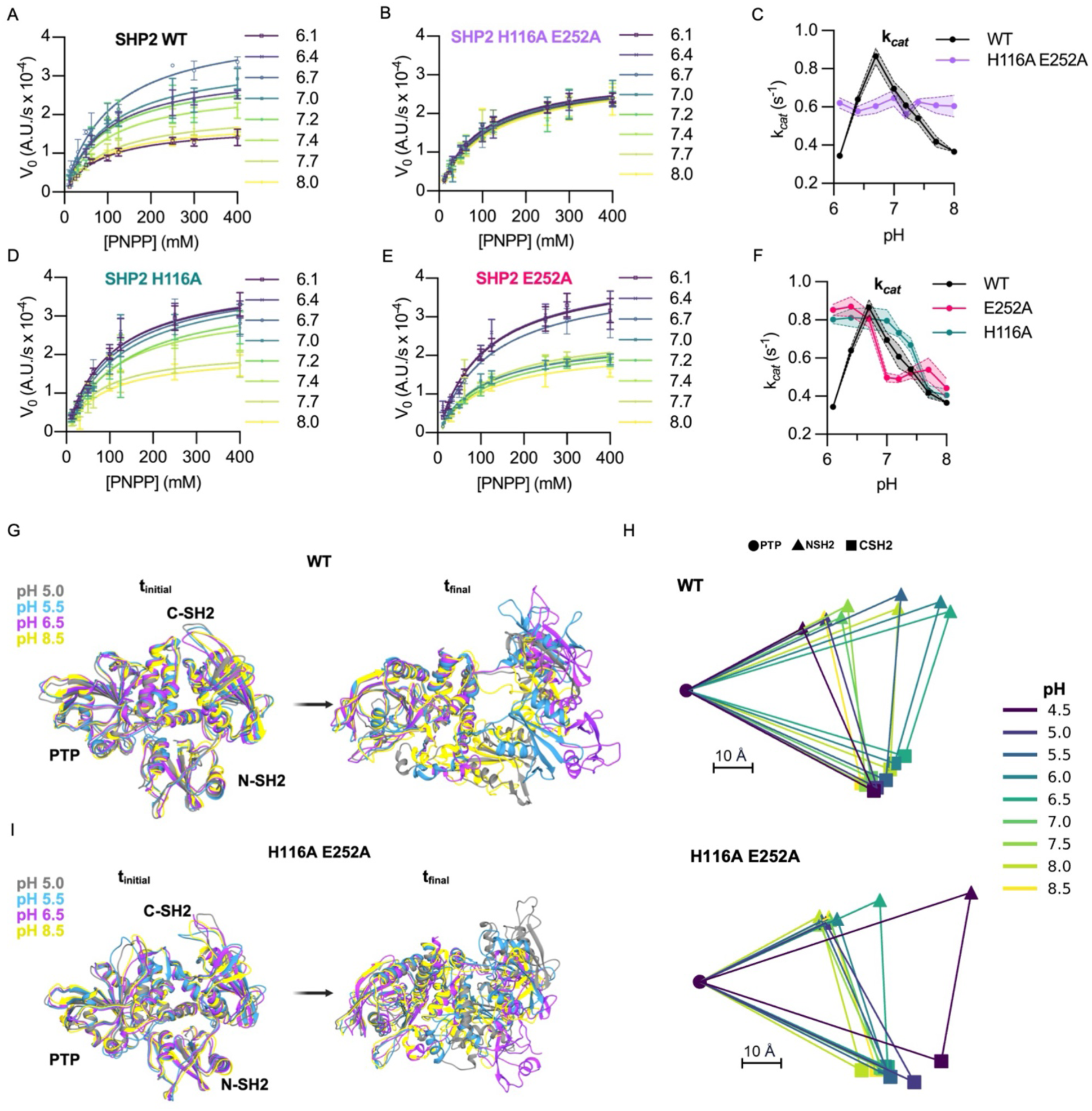
Molecular mechanism of SHP2 pH sensing requires both His116 and Glu252. (A) Wild-type (WT) SHP2 in vitro phosphatase activity curves with increasing concentrations of generic substrate p-Nitrophenyl Phosphate (PNPP) at buffer pH ranging from 6.1 to 8.0. (mean ± SEM; N=3 from ≥2 different protein preparations) (B) Double mutant (H116A E252A SHP2) in vitro phosphatase activity, assays performed as in A. (mean ± SEM; N=3 from ≥2 different protein preparations) (C) Plot of k*_cat_* vs. pH for WT and double mutant (H116A E252A SHP2). Calculated from activity curves in A and B. (mean ± SEM) (D-E) Single-mutant in vitro phosphatase activity for (D) H116A-SHP2 and (E) E252A-SHP2. Assays performed as in A. (mean ± SEM; N=3 from ≥2 different protein preparations) (F) Plot of K*_cat_* vs. pH for WT and single mutant H116A SHP2 and E252A SHP2. Calculated from activity curves in A, D and E. (mean ± SEM) (G) CpHMD was performed on WT SHP2 at pH values from 4.0-10.0. Shown are overlapping views of WT SHP2 structures at the start of CpHMD simulation (t = 0 ns) and end of simulation (t = 8 ns) for pH values 5.0 (grey), 5.5 (cyan), 6.5 (magenta) and 8.5 (yellow). (H) Average Interdomain center of mass distances for WT and H116A E252A SHP2 for the last 3 ns of the simulation (I) CpHMD was performed on H116A E252A SHP2 at pH values from 4.0-10.0. Shown are overlapping views of H116A E252A SHP2 structures at the start of CpHMD simulation (t = 0 ns) and end of simulation (t = 8 ns) for pH values 5.0 (grey), 5.5 (cyan), 6.5 (magenta) and 8.5 (yellow).

Reactions with bovine serum albumin (BSA) addition showed no pH-dependence, confirming that buffer pH is not independently affecting non-enzymatic PNPP hydrolysis in these assays (**Fig. S3D**). To test whether the identified network is responsible for SHP2 pH-dependent activity, we generated a SHP2 double mutant where both H116 and E252 were mutated to non-titratable alanine residues. We repeated the phosphatase assay and found that the activity of SHP2-H116A/E252A was pH-independent (**Fig. 2B**). While k*_cat_* of SHP2-H116A/E252A was reduced compared to WT SHP2, there was no pH-dependence of k*_cat_*, K_m,_ or catalytic efficiency (**Fig. 2B, C, S3A, S3C)**. This demonstrates that H116 and E252 are required for pH-dependent activity of SHP2.

Our in silico pK_a_ prediction analysis suggests that H116 and E252 function within networks of conserved ionizable residues. To probe this interaction network in more detail, we next generated the single point mutants (SHP2-H116A and SHP2-E252A) and found that both mutants retained pH-dependent phosphatase activity, but each with a single apparent pK_a_ (**Fig. 2D-F**). For both mutants, k*_cat_* was maximal at low buffer pH and reduced with increasing pH (**Fig. 2F, S3A, S3C**). We predicted that the pH-dependent activity observed in the single point mutants reflects the pK_a_ of the remaining titratable residue. We compared apparent pK_a_s of the mutants and found that SHP2-H116A mutant had an apparent pK_a_ of 7.5 (**Fig. 2F, S3A, S3E**), which correlates with the higher apparent (pK_a2_) of wild-type SHP2 (∼7.1) and the in silico predicted pK_a_ of E252 (∼7.4) (**Fig. 1D**). The E252A mutant has a lower apparent pK_a_ of 6.8 (**Fig. 2F, S3A, S3E**), which matches the lower apparent (pK_a1_) of the wild-type SHP2 (∼6.7), and the in silico predicted pK_a_ of His116 (6.8) (**Fig. 1D).** This suggests that in the H116A point mutant, E252 drives the observed pH-dependent activity while in the E252A point mutant, H116 drives the observed pH-dependent activity. Compared to the wild-type SHP2 activity profiles, this suggests that H116 titration regulates activity between pH 6.1 to 7.0 while E252 titration regulates activity between pH 7.0 to 8.0. Thus, Glu252 and His116 function cooperatively to confer wild-type SHP2 pH-dependent activity.

Because E249 interacts with both Glu252 and His116, we next explored the role this bridging glutamate plays in the pH sensing mechanism of SHP2. Interestingly, we observed similar pH-dependent activity profiles of SHP2-E249A compared to WT (**Fig. S2A, F**). The E249A mutant was still pH sensitive with biphasic k*_cat_* and catalytic efficiency pH profiles (**Fig. S3C, S3G**). However, the apparent pK_a1_ and pK_a2_ of the E249A mutant were both shifted higher (6.8 and 7.7) when compared to wild-type SHP2 (6.7 and 7.1) (**Fig. S3A, S3C**). This suggests that E249 is not acting as a direct proton conduit or relay between the identified pH sensing residues (H116 and E252) but instead the presence of E249 tunes the pK_a_ values of these residues through the interaction network.

### CpHMD reveals pH-dependent residue conformations and SH2 domain release in SHP2

Due to the clustering of pH-sensitive residues at the SH2 and PTP domain interface, we hypothesized that pH-dependent binding of the inhibitory SH2 domains drives pH-dependent SHP2 activity. We used continuous constant pH molecular dynamics (CpHMD) to probe pH-dependent structural conformations of wild-type SHP2. In classical molecular dynamics (MD) simulations, solvent pH is considered implicitly by fixing protonation states based on empirical or predicted residue pK_a_ values. However, protein conformational dynamics may depend explicitly on residue protonation states, so conventional MD methods cannot capture how dynamic changes in residue protonation affect structural dynamics. CpHMD addresses this gap by sampling protonation and conformation states simultaneously(*34*) and lends insight into proton-coupled mechanisms that are not captured in conventional MD methods.

We performed CpHMD simulations of SHP2 using the GBNeck2 implicit solvent model for pH values ranging from 4.0 to 10 (in increments of 0.5) (based on(*34*), see methods; trajectories in **Movies 1-13**). Each simulation was run for 8 nanoseconds, allowing for convergence of backbone RMSD and protonation states, for an aggregate 104 nanoseconds of total simulation time. In agreement with the in vitro results, the pK_a_ values calculated from SHP2 CpHMD showed that H116 had a lower pK_a_ (6.0) compared to E252 (8.4) (**Fig. S4A, S4B**). In addition, we observed and quantified coordinated pH-dependent changes in the side chain conformations of H116 and E252. (**Fig. S4C-D**). At low (4.5) and high (8.5) pH, H116 and E252 are pointed toward each other, but at optimal pH for activity (6.5), E252 flips out and H116 swings in, disrupting its local helix (**Fig. S4C-D**). From the CpHMD trajectories, we noted that the SH2 domain remains rigid and tightly bound to the PTP domain at pH values less than 5.5 and higher than 8.0 (**Fig. 2G-H**, **Movie4 (2SHP_pH5.5) and Movie9 (2SHP_pH8.0)**).

However, at optimal activity pH (6.5) the SH2 domain is more dynamic and loosely bound to the PTP domain (**Fig. 2G-H, Movie6 (2SHP_pH6.5**)). To assess interdomain conformations, we quantified the center of mass distances between the N-SH2 or C-SH2 domain and the PTP domain during the final 3 nanoseconds of the trajectories. We found distinct pH-dependent dynamics between the N-SH2 and PTP domain, with maximal interdomain distances (65Å) at pH 6.5 and minimal distances (30Å and 35Å) at pH 4.5 and 8.5 respectively (**Fig. 2H, S4E)**. We observed that dynamics between the C-SH2 and PTP domain followed the same pH-dependent trend, with maximal distances (54Å) achieved at pH 6.5 **(Fig. 2H, S4F).** While interdomain orientations were not included in this analysis, the interdomain distance trends alone suggest pH-dependent SH2 domain release and are in striking agreement with pH-dependent in vitro WT SHP2 activity experiments.

We also performed CpHMD simulations for the H116A E252A SHP2 mutant using the same protocol as for WT to determine if these mutations also abrogate the pH dependent interdomain dynamics in silico. Encouragingly, we observed that double mutant exhibited a more closed domain conformation when compared to WT across the pH range 6.0-8.0 (**Fig. 2 H-I, S5A-C**). Importantly, the double mutant did not exhibit the strong pH-dependent inter-domain dynamics that we observed with WT SHP2 (**Fig. 2H-I**). Our CpHMD results indicate a loss of pH-dependent interdomain dynamics in the mutant protein, which is in agreement with the in vitro activity results. Taken together, our CpHMD data support a molecular mechanism by which WT SHP2 has pH-dependent SH2 domain binding, with His116 and E252 being required for pH-dependent SH2 domain conformational dynamics.

### Intracellular pH regulates SHP2 activity and downstream signaling in MCF10A cells

We next determined whether SHP2 activity is regulated by physiological pH changes in cells. We first experimentally manipulated intracellular pH (pHi) in normal breast epithelial cells (MCF10A), using inhibitors of the sodium proton exchanger (EIPA) and sodium bicarbonate transporter (S0859) to lower pHi from 7.45±0.12 to 7.10±0.01 and 30 mM ammonium chloride to raise pHi to 8.00±0.27 (**Fig. 3A**). After one hour of pHi manipulation, we assayed for endogenous SHP2 basal activity by immunoblotting for phosphorylation of SHP2 at tyrosine 542 (Y542), which is required for activity(*35*). We found that endogenous SHP2 activity is pH sensitive, with increased pY542 levels at low pHi and decreased activity at high pHi compared to control (**Fig. 3B, C**). To investigate downstream signal transduction, we measured SHP2 binding to its signaling-active binding partner, Gab1(*36*) using a co-immunoprecipitation assay. SHP2 immune complexes contained significantly more Gab1 when pHi was low compared to control and high pHi (**Fig. 3B, D**). This demonstrates that endogenous SHP2 has pH-dependent activity in MCF10A cells, with higher activity and binding to the Gab1 signaling partner at low pHi compared to control and high pHi.

**Figure 3.**
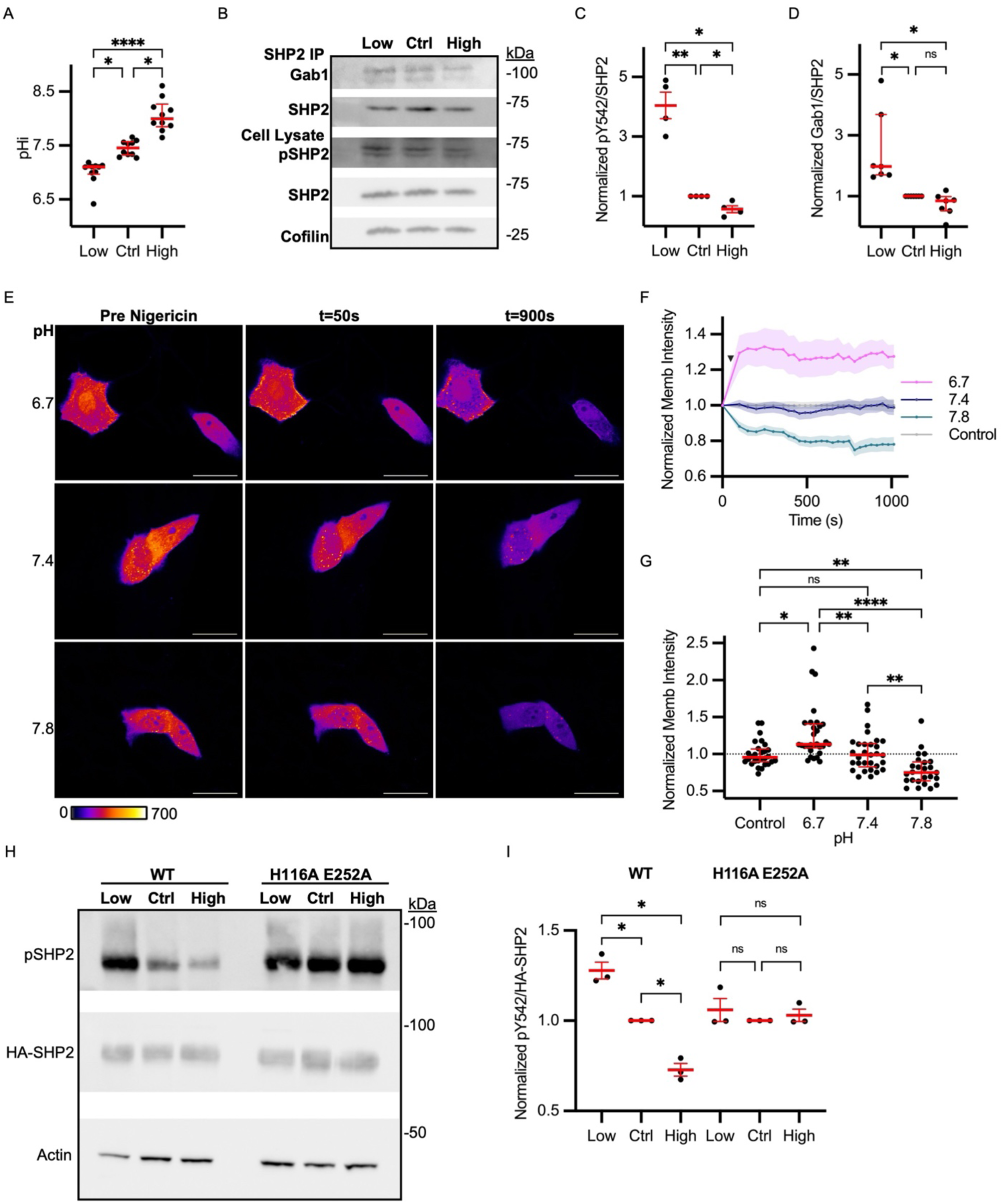
Intracellular pH regulates SHP2 activity and downstream signaling in MCF10A cells. (A) pHi measurements of MCF10A cells. Cells were treated with 25 μM EIPA + 30 μM S0858 for 1 hour to lower pH to 7.10. To raise pHi, cells were treated with 30 mM ammonium chloride for 1 hour to raise pH to 8.00. Untreated cells had a pHi of 7.45. Scatter plot shows (median ± interquartile range, N =10) (B) Representative immunoblots of SHP2, Gab1, and phospho-SHP2 (pY542) from SHP2 immune complexes (SHP2 IP) or whole cell lysates (Cell Lysate) isolated from MCF10A cells prepared as in A. (C-D) Quantification of phospho-SHP2 (C) and Gab1 Co-IP (D) for replicate data collected as in B. Data was normalized to control in each biological replicate. Scatter plot shows (mean ± SEM, N=4) in C and (median ± IQR, N=7) in D. (E) Representative images of MCF10A cells expressing the SHP2 activity reporter (Grb2 TagBFP) pseudocolored on an intensiometric scale. Images show cells prior to manipulating pHi with nigericin buffer (Pre-Nigericin) (see methods for details), 50s after manipulating pHi, and 900s after manipulating pHi. Scale bars: 25μm (F) Quantification of images collected as in E. Membrane intensity of SHP2 activity reporter over time was photobleach-corrected and then normalized to initial intensity. Line trace shows (mean ± SEM) from single-cell data (6.7 pH, n=30, 7.4 pH, n=30, 7.8 pH, n=25, control, n=28) collected across N=3 biological replicates. (G) Quantification of endpoint membrane intensities of single cells collected as described in F. Scatter plot shows (median ± interquartile range, N = 5) (H) Representative immunoblot of lysates prepared from MCF10A cells expressing either WT SHP2 or H116A E252A SHP2. Immunoblots show total and pY542-SHP2 under low, control, and high pHi conditions. (I) Quantification of replicate data collected as in H. Scatter plots show (mean ± SEM, N=3). Intensities were normalized to the corresponding control condition. For A and G significance was determined using the Kruskal-Wallis test. For C and I significance was determined using a ratio paired t-test to compare between treatment conditions and a one-sample t-test to compare to control. For D significance was determined using a Wilcoxon test to compare between treatment conditions and a Wilcoxon sign-ranked test to compare to control. * p<0.05, **p<0.01, ***p<0.001, ****p<0.0001.

We next confirmed the effect of pHi dynamics on basal activity of endogenous SHP2 using a live, single-cell reporter of SHP2 activity (tagBFP-Grb2)(*37*). We validated that the tagBFP-Grb2 reporter was robustly recruited to the cell membrane upon EGF stimulation (active SHP2) and that this recruitment was abolished by both a specific SHP2 inhibitor (PHPS)(*38*) and an upstream EGFR inhibitor (Erlotinib) (**Fig. S6A**). We also validated inhibitor efficacy and specificity using western blots to assess EGFR and SHP2 phosphorylation (**Fig. S6B**). Erlotinib treatment reduced EGF-induced pEGFR (pY1068; Grb2 phosphosite) and pSHP2 (pY542), whereas PHPS reduced pSHP2 while leaving pEGFR levels unaffected (**Fig. S6B**). These data show that membrane recruitment of the biosensor is dependent on SHP2 activity.

After validating tagBFP-GRB2 as a specific SHP2 activity biosensor, we next measured biosensor recruitment to the MCF10A cell membrane under basal (unstimulated) conditions with and without pHi manipulation using the protonophore nigericin (see methods for details). When the pHi of the MCF10A cells was pinned to 6.7, we observed increased biosensor recruitment to the cell membrane, indicating increased SHP2 activity (**Fig. 3E-G).** Conversely, we observed no change in biosensor recruitment when MCF10A cell pH was pinned to basal pHi conditions (7.4) and reduced biosensor recruitment when the pHi of MCF10A cells was pinned at 7.8 (**Fig. 3F-G)**. Importantly, initial biosensor intensity at the membrane was not different prior to nigericin-based pHi manipulation (**Fig. S6C**) and increased biosensor recruitment at low pHi was abolished with the addition of PHPS (**Fig. S6D**). This shows that endogenous SHP2 basal activity is sensitive to pHi dynamics in single living cells.

To confirm that the pH sensing network of H116 and E252 mediates pH-dependent SHP2 activity in cells, we overexpressed HA-tagged WT SHP2 and SHP2-H116A/E252A in MCF10A cells and quantified SHP2-pY542 levels with and without pHi manipulation. We found that overexpressed WT SHP2 recapitulated the endogenous SHP2 results, with increased activity (SHP2-pY542) at low pHi and decreased activity at high pHi compared to control MCF10A (**Fig. 3H, I**). As expected, the double mutant SHP2-H116A/E252A had pH-independent activity (SHP2-pY524) in MCF10A (**Fig. 3H, I**), confirming that His116 and E252 are required mediators of pH-sensitive SHP2 activity. Through our cell-based analyses, we show that SHP2 activity is sensitive to physiological changes in pHi, with increased basal activity at low pHi, and that H116 and E252 mediate pH-dependent activity of SHP2.

### pH sensing nodes cluster at SH2 interface in other SH2 signaling proteins

SH2 domains are modular signaling domains that bind to phosphotyrosine residues, which have experimentally determined pK_a_s that are close to physiological (5.7-6.1)(*39*, *40*) and are highly structurally and sequence-conserved across signaling proteins(*29*). We hypothesized that networks of residues with physiological pK_a_s serve a functional role in regulating pH-dependent SH2 binding to phosphotyrosine residues across SH2 domain-containing proteins. To explore this, we applied our in silico computational pipeline to a collection of SH2 domain-containing proteins with multiple solved structures. We found that 90% of SH2 domain-containing proteins tested had ionizable residue networks with pK_a_s predicted to be shifted into the physiological pH range (**Fig. S7**). As we observed with SHP2, these ionizable residue networks clustered at the SH2 binding interfaces (**Fig. S7**). Interestingly, we observed this pattern of clustering across functional protein classes—finding it in phosphatases (SHP1) (**Fig. S7A**), kinases (ZAP70, JAK1 JAK2, and ABL1) (**Fig. S7B-D, H**), transcription factors (STAT6, STAT5A) (**Fig. S7E-F**), and GTPase activating proteins (CHIO) (**Fig. S7G**).

As a control, we also analyzed a collection of SH3 domain-containing signaling proteins. SH3 domains are modular signaling domains that are similarly highly evolutionarily and structurally conserved(*41*), but bind proline-kink regions(*42*) where electrostatics should not drive function. While we identified ionizable networks with physiological pK_a_s in some SH3 domain-containing proteins, the shifted residues did not cluster at the binding interfaces of SH3 domains (**Fig. S7H, S7I-K**). This analysis suggests that ionizable networks with upshifted pK_a_s are not a general feature of all signaling domain interaction interfaces but instead are enriched at functional SH2 domain interfaces.

### Analysis pipeline applied to Src identifies four networked ionizable residues with predicted physiological pK_a_s

We next selected one of the hits from the screen of SH2 domain-containing signaling proteins (Src) to further validate the role of pH in the allosteric regulation of SH2 domain-containing signaling proteins. The signaling kinase Src has three globular domains: a kinase domain (SH1), an inhibitory SH2 domain that binds phosphotyrosines, and an inhibitory SH3 domain that binds proline-rich regions (**Fig. 4A**). The SH2 domain inhibits kinase activity when bound intramolecularly to the C-terminal phosphotyrosine (pY527)(*42*). However, when the SH2 and SH3 domains are released, pY527 is dephosphorylated, the activation loop unfurls, and the SH1(kinase) domain autophosphorylates a conserved tyrosine (Y416) in the activation loop (A-loop). Activation loop unfolding and phosphorylation are required for full activation and facilitate Src binding to downstream targets(*42*).

**Figure 4:**
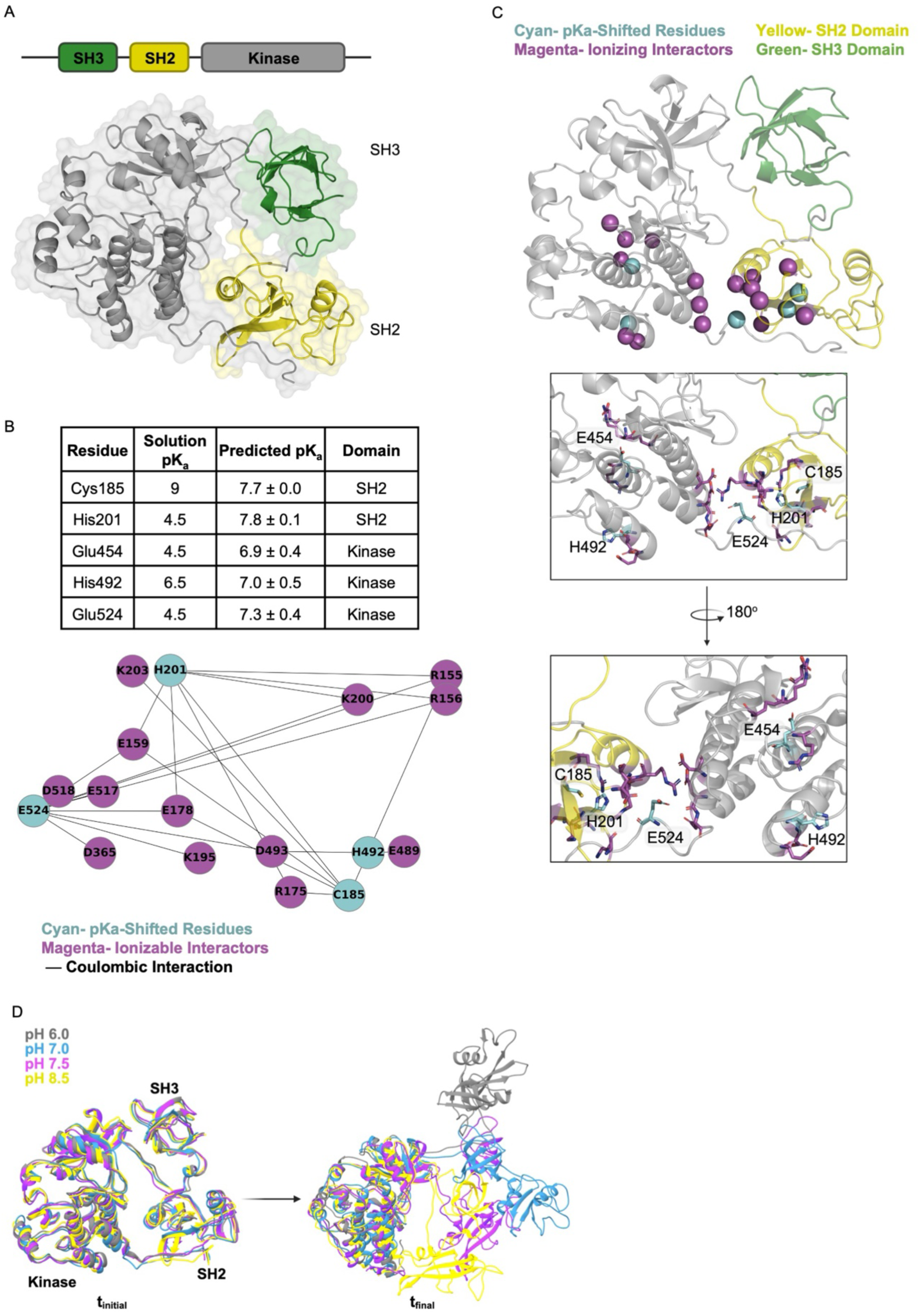
Residues with predicted physiological pK_a_ shifts cluster at the binding interface of the SH2 and kinase domains in Src. (A) Crystal structure of pilot protein Src shown in cartoon and surface format (PDB ID:2SRC). Kinase domain colored in grey, SH2 domain colored in yellow, and SH3 colored in green (B) Table of predicted pK_a_s for shifted residues (mean ± SD). Residue interaction network of residues with predicted pK_a_ shifts (cyan) and their coulombic ionizable interactors (magenta). Edge length reflects the strength of the interaction, with stronger coulombic interactions having shorter edge lengths. (C) Structure of Src (PDB ID:2SRC) showing the kinase domain in grey, SH2 domain in yellow, and SH3 domain in green. Residues identified through in silico ionizable network prediction pipeline shown in spheres. Residues with predicted pK_a_ shifts (cyan) cluster with ionizable interactors (magenta) across the kinase-SH2 domain interaction interface of Src. Zoom in shows Src structure at the kinase-SH2 domain interaction interface. Networked residues from B and C are shown in stick: residues with predicted pK_a_ shifts in cyan, interactors in magenta. (D) CpHMD (see methods for details) was performed on Src at pH values from 4.0-10.0. Shown are overlapping views of Src structures at the start of CpHMD simulation (t = 0 ns) and end of simulation (t = 8 ns) for pH values 6.0 (grey), 7.0 (cyan), 7.5 (magenta) and 8.5 (yellow).

We processed seven structures of Src available in the PDB through the computational pipeline. Our analysis identified 5 residues predicted to have physiological pK_a_s: C185 (pK_a_ of 7.7 ± 0.0), H201 (pK_a_ of 7.8 ± 0.1), E454 (pK_a_ of 6.9 ± 0.4), H492 (pK_a_ of 7.0 ± 0.5), and E524 (pK_a_ of 7.3 ± 0.4) (**Fig. 4B, C)**. Four residues (C185, H201, E524, and H492) are networked in a cluster at the interface between the SH2 domain and the kinase domain (**Fig. 4B, C**). E524 is in the C-terminal tail of Src and electrostatically interacts with H201 and C185 in the SH2 domain through bridging glutamate residues (E159, E178) and interacts with H492 in the kinase domain through a bridging aspartate residue (D493) (**Fig. 4B, C**). While E454 has a predicted physiological pK_a_, it is in the kinase catalytic site and distant from the SH2 regulatory domain interface (**Fig. 4B**). This analysis of Src again demonstrates that ionizable residue networks with predicted physiological pK_a_s cluster at SH2 but not SH3 domain interfaces.

We determined the structural conservation of the identified residues in Src and found very high structural conservation of the ionizable network region and individual residues (**Fig. S8A-C**). The COSMIC database(*43*) also showed recurrent somatic mutations at each of these sites: C185, H201, H492, and E524 (**Fig. S8D**). Recurrent charge-changing somatic mutations were found in H201, H492, and E524 (Fig S8D, E), suggesting functional importance of the identified network and potential for pH-sensitive network disruption in cancer.

### CpHMD reveals pH-dependent residue conformations and SH2 domain release in Src

We next assessed pH-dependent structural dynamics in Src using CpHMD (see methods; trajectories in **Movies 14-26**). While we were not able to generate a pK_a_ for C185 using the CpHMD trajectories (**Fig. S9A**), all other identified residues had upshifted and near-physiological pK_a_s calculated from the CpHMD trajectories: (H201, pK_a_ of 7.3; E454 pK_a_ of 8.2; H492 pK_a_ of 7.0; E524 pK_a_ of 6.8) (**Fig. S9B-E**). Local side chain movement showed pH-dependent changes in the identified residue network, suggesting pH-dependent interaction pairs (**Fig S9F, G**). Dihedral angles change with pH for all identified residues, with coupled dynamics observed for connected residues (H201, C185, and H492) between pH 6.5 and 7.5 while E524 dihedral angles change at pH values above 7.0 **(Fig S9F**). The CpHMD trajectories also show pH-dependent changes in interdomain interactions between the Src kinase domain and the inhibitory SH2 and SH3 domains (**Fig. 4D**). In agreement with the SHP2 dynamics, the SH2 domain of Src was bound more tightly to the kinase domain at high pH (8.5) compared to low pH (6.0), where both the SH3 and SH2 domain were unbound (**Fig. 4D, Movie18 (2SRC_pH6.0), Movie23 (2SRC_pH8.5)**). We also observed intermediate domain conformations in the pH 7.0 and pH 7.5 trajectories, with Src adopting a conformation where the SH3 domain binds tightly while the SH2 domain is more dynamic (**Fig 4D, Movie20 (2SRC_pH7.0), Movie21 (2SRC_pH7.5**)). To assess interdomain conformations, we quantified interdomain distances between either the SH2 or SH3 domain and the central kinase domain of Src during the final 3ns of the trajectories. We found pH-dependent SH2-kinase interdomain distances with a maximum (least binding) at pH 6.0 and minimums (tight binding) at high (8.5) and low (4.5) pH (**Fig. S9H)**. We observed similar pH-dependent trends in pH-dependent SH3-kinase interdomain distances, with a maximum (least binding) distance at pH 5.5 and minimums at high (8.5) and low (5.0) pH (**Fig. S9I**). To rule out general pH-dependent structural compaction in these CpHMD trajectories, we also assessed SH3-SH2 domain distances and found no pH dependence (**Fig. S9J**). While these short trajectories are less likely to capture large protein domain movements, the trends across the SHP2 and Src CpHMD analyses are strikingly consistent: low physiological pH (6.0-7.0) leads to SH2 release from the catalytic domain compared to high physiological pH (7.0-8.0) where SH2-catalytic interdomain interactions are increased.

### Intracellular pH regulates Src activity and signaling in MCF10A cells

We next determined whether basal kinase activity of endogenous Src is sensitive to physiological pH changes in normal epithelial cells. We assessed Src activity by immunoblot quantification of the levels of pY527, a proxy for inhibited Src, and pY416, a proxy for active Src (**Fig. 5A**). We found that endogenous Src activity was pH sensitive, with increased activity at low pHi compared to control and high pHi (**Fig. 5B-D**) with no change to total endogenous Src levels (**Fig. S10A**). Importantly, the pH-dependent trends in pY527 and pY416 both support the hypothesis that low pHi leads to SH2 domain release, loss of the inhibited Src phosphosite (pY527) (**Fig. 5C**), and gain of the active Src phosphorylation site (pY416) (**Fig. 5D**). Next, we performed immunofluorescence labeling and single-cell analysis of Src activity in MCF10A cells and found that active Src (pY416) was increased at low pHi and decreased at high pHi compared to control cells (**Fig. 5E-G, S10B, C**). We stimulated cells with EGF to determine how pHi regulates Src in the context of growth factor stimulation. We observed significant increases in Src activation (pY416) in EGF stimulated control cells, demonstrating that growth factor stimulation can drive increased Src activation (**Fig. 5E-G**). Interestingly, there was no difference in pY416 levels at low pHi compared to control cells stimulated with EGF, suggesting low pHi phenocopies Src activation via growth-factor stimulation (**Fig. 5E-G**). However, when high pHi cells were stimulated with EGF, activation of Src was lower compared to control cells both with and without EGF stimulation (**Fig. 5E-G, S10B, C**). This indicates that high pH inhibition of Src is sufficient to attenuate the growth factor receptor activation of endogenous Src.

**Figure 5.**
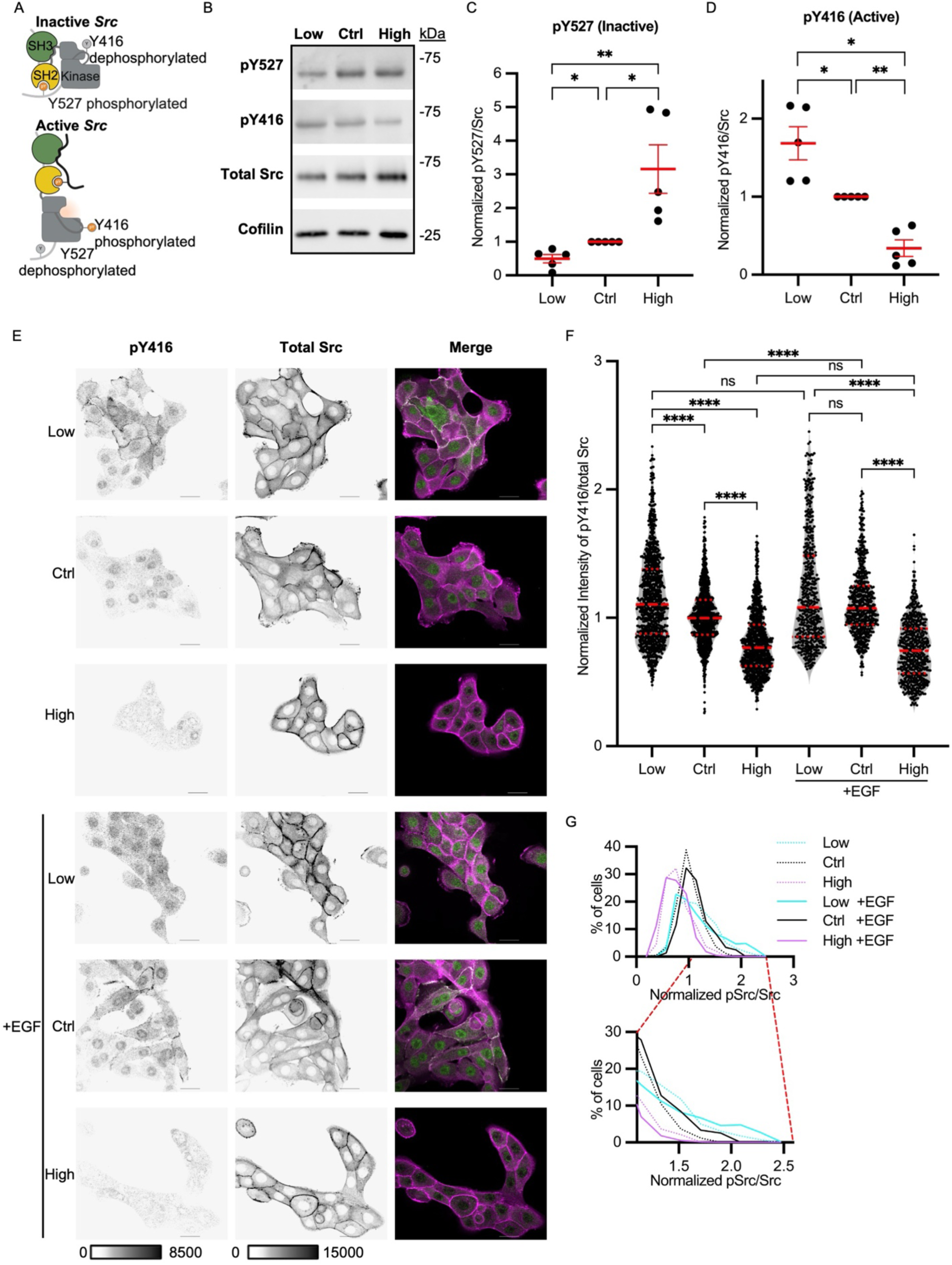
Intracellular pH regulates Src activity and localization in MCF10A cells. (A) Cartoon of Src activation (B) Representative immunoblot from MCF10A lysates prepared from cells maintained for 1 hour at low or high pHi or untreated (control). Immunoblots show total Src and indicated Src phosphorylation sites that reflect protein activation pY527 (inactive) and pY416 (active). (C-D) Quantification of pY527 (inactive) (C) and pY416 (active) (D) Src from replicate data collected as in B. Data was normalized to control in each biological replicate. Scatter plots show (mean ± SEM, N=5) (E) Representative confocal images of MCF10A cells fixed and stained for total Src and pY416 (active) Src at given pHi with and without EGF stimulation (10 ng/mL for 5 minutes). Total Src and pY416 are shown in inverse mono intensity display. In the merged overlay, total Src is shown in magenta and pSrc is shown in green, areas of high co-localization show white. Scale bar: 25μm. (F) Quantification of single-cell data collected as in E. Levels of pY416 Src are normalized to total Src in each single cell and to control for each biological replicate. Violin plot shows individual cells as points and (median ± IQR, Low, n=984, Control, n=938, High, n=903, Low + EGF, n=525, Control + EGF, n=595, High + EGF, n=599) collected from 6 biological replicates for - EGF conditions and 3 biological replicates for + EGF conditions. (G). Histogram and zoomed in view of single cell data for data shown in F. Low pHi treatment conditions shown in cyan, control in black, and high pHi treatment shown in magenta. For C and D, significance was determined using a ratio paired t-test to compare between treatment conditions and a one-sample t-test to compare to control. For F, significance was determined using the Kruskal-Wallis test. * p<0.05, **p<0.01, ***p<0.001, ****p<0.0001.

It was at first surprising that growth factor stimulation at low and high pHi did not further increase Src activation (pY416) compared to unstimulated cells with matched pHi (**Fig. 5F**).

However, when comparing pY416 Src distributions, we did observe statistically significant differences between EGF stimulated and unstimulated cells across all pHi conditions (**Fig. S10B, C**). Our data suggest that pHi manipulation can modulate Src activity in the context of growth factor signaling as well as in basal state.

To determine cellular outcomes of pHi-dependent Src activation, we next assessed downstream signal transduction by monitoring downstream activation of ERK (pT202/ pY204) and E-cadherin (pY685). In both cases, pathway activation downstream of Src recapitulated the pH-dependent activity trends, with increased activation at low pHi and decreased activation at high pHi compared to control cells (**Fig. S10D-G**). In control cells, EGF stimulation produced statistically significant increases in downstream activation of ERK (pT202/ pY204) and E-cadherin (pY685) (Fig S10D-G). With EGF stimulation, there was no difference in downstream pathway activation at low pHi compared to control, again suggesting maximal activity is achieved (**Fig. S10D-G**). Finally, high pHi produced lower downstream activation both with and without EGF stimulation compared to matched control conditions (**Fig. S10D-G**). This indicates that low pHi phenocopies EGF stimulation of control cells while high pHi inhibition of Src can attenuate pathway activation even in the presence of extracellular growth factor.

To determine whether the identified residue network in Src mediates pH-dependent activity, we overexpressed either FLAG-tagged WT-Src or the quadruple mutant Src-C185A/E454A/H492A/E524A in MCF10A cells. We then quantified pY527 and pY416 levels with and without pHi manipulation by immunoblot. We found that overexpressing WT Src recapitulated the pH-dependent endogenous Src activity results, with decreased phosphorylation of the inhibitory phosphosite (pY527) at low pHi and increased pY527 at high pHi compared to control MCF10A (**Fig. 6A, B**). As with endogenous Src, we also observed pHi-dependent phosphorylation of pY416, with increased active Src (pY416) at low pHi compared to control and high pHi (**Fig. 6A, C)**. Taken together, these data demonstrate that pHi allosterically regulates Src inside cells, with low pHi promoting the active state (pY416, Y527) while high pHi promotes a more inhibited state (Y416, pY527). As expected, the quadruple Src mutant had pH-independent phosphorylation of both pY416 and pY527 while maintaining measurable catalytic activity (**Fig. 6A-C**). Our cell-based analyses show that Src activity is sensitive to physiological pHi changes, with increased activity at low pHi, and that the identified ionizable residue cluster is required for pH-dependent Src activity.

**Figure 6.**
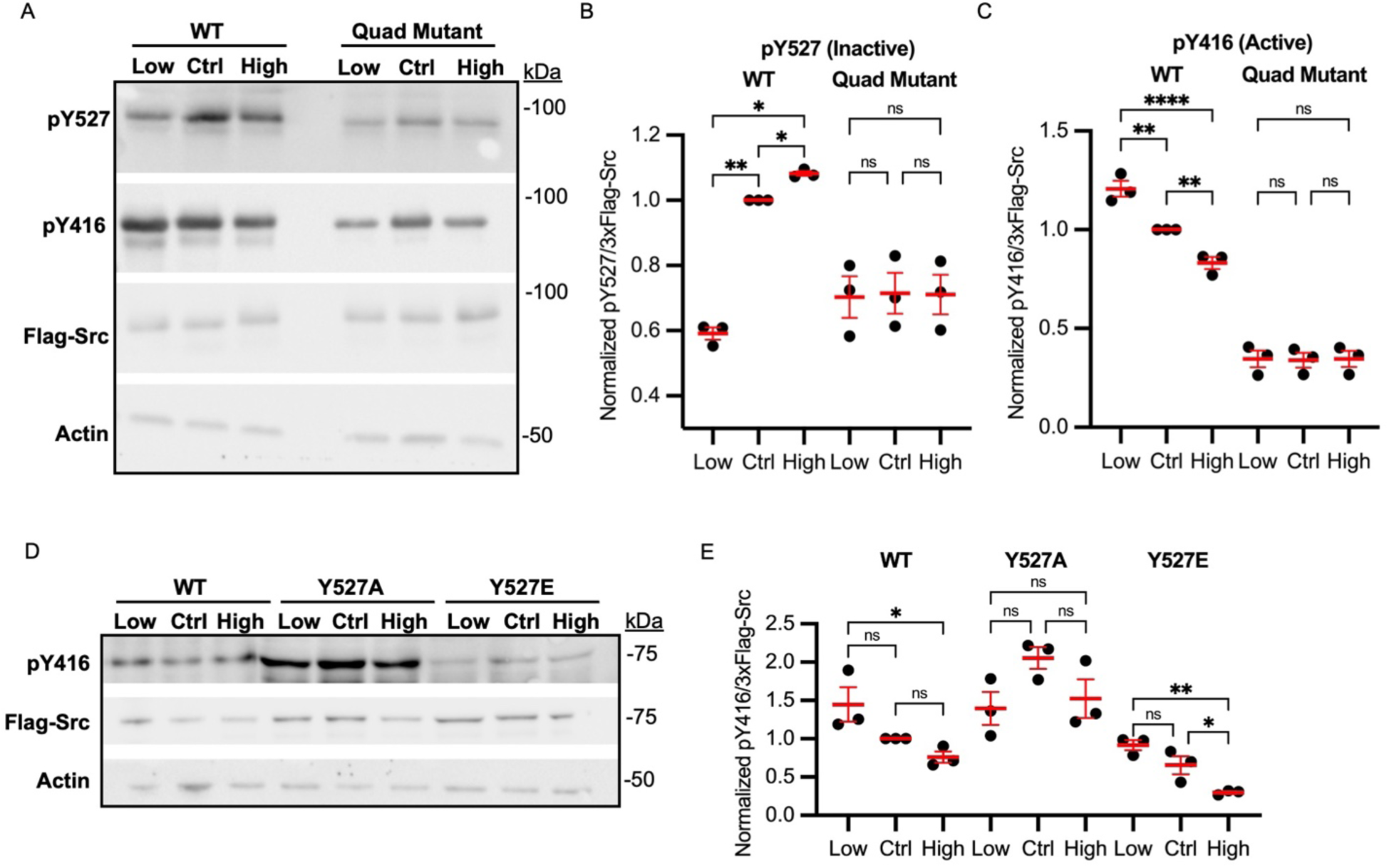
Src pH-dependent activity requires an intact ionizable residue network (C185 H201 H492 E524) (A) Representative immunoblot from MCF10A lysates expressing either WT Src or the Quad Mutant (Src-C185A/H201A/H492A/E524A) and maintained for 1 hour at low or high pHi or untreated (control). Immunoblots show total Src and indicated Src phosphorylation sites that reflect protein activation pY527 (inhibited) and pY416 (active). (B) Quantification of pY527 (inactive) from replicate data collected as in g. Data was normalized to control in each biological replicate. Scatter plots show (mean ± SEM, N=3) (C) Quantification of pY416 (active) from replicate data collected as in g. Data was normalized to control for each biological replicate. Scatter plots show (mean ± SEM, N=3) (D) Representative immunoblot from MCF10A lysates expressing either WT Src, Y527A Src, or Y527E Src and maintained for 1 hour at low or high pHi or untreated (control). Immunoblots show total Src and indicated active Src phosphorylation site (pY416). (E) Quantification of pY416 (active) from replicate data collected as in D. Data was normalized to WT control for each biological replicate. For B and C, significance was determined using a ratio paired t-test to compare between treatment conditions and a one-sample t-test to compare to WT control. For E, significance was determined using a one-way ANOVA and a one-sample t-test to compare to WT control. * p<0.05, **p<0.01, ***p<0.001, ****p<0.0001.

To further probe contributions of individual residues in the identified residue network to pH-sensitive Src activity, we generated single point mutants (Src-C185A, Src-E524A, Src-H201A, and Src-H496A) and measured their activity in MCF10A cells. First, we found that the E524A mutant pushed Src into a more active, but still pH-dependent, state: Y527 was significantly dephosphorylated compared to WT Src and pH-independent while Y416 phosphorylation was still pH-dependent but increased compared to WT (**Fig. S11A-C**). This suggests E524 is required for the observed pH-dependence of pY527 in wild-type Src: without E524, affinity for pY527 is reduced and the Src C-terminal tail release from the SH2 domain is pH-insensitive in the physiological range. We can also conclude from the E524A data that the remaining residue network (C185, H201, and H495) is sufficient to preserve pH-dependent phosphorylation of active-loop residue (Y416) (**Fig. S11A-C**).

We found that C185A and H496A still exhibited trends of pH-dependent phosphorylation of both Y527 and Y416 (**Fig. S11A-F**), suggesting C185 and H496 are not essential for either pH-sensitive phosphorylation event on Src. However, because we cannot easily titrate across wider pHi ranges in these cells (e.g. 6.1-8.0), we cannot rule out the possibility that C185 and H496 may be critically important for tuning the pKa of the entire residue network. Finally, the H201A mutant locked Src into an inhibited and pHi-independent conformation, with significantly reduced Y416 phosphorylation compared to WT (**Fig. S11D-F**). This suggests H201 plays a predominant role in regulating the pH-dependent active-loop phosphorylation of Y416 observed in both WT and Src-E524A.

Finally, to determine the specific role pH plays in regulating the well-characterized (*44*, *45*)phospho-tyrosine activation of Src, we assessed pH-dependent Src activity with Y527 phosphomimetic mutants. We mutated Y527 to a non-titrating alanine that mimics a constitutively dephosphorylated Y527 and to a titratable glutamate that mimics a constitutively phosphorylated Y527. As expected, Src-Y527A was constitutively active (high pY416) and pH-independent (**Fig. 6D,E**). This data phenocopies Src constitutive activation when Y527 is mutated or deleted (*44*). Conversely, we found that Src-Y527E activation was still pH dependent, with higher activity at low pHi and lower activity at high pHi when compared to control (**Fig. 6D,E**). Taken together, these data suggest phosphorylation of Y527 contributes to the modulation of pH-dependent wild-type Src activation. Thus, our data suggest pHi dynamics function as a rheostat to modulate WT Src activity in concert with traditional modes of Src regulation (phosphorylation).

## Discussion

Our data show that our in silico analysis pipeline can be used to predict and mechanistically probe pH-dependent protein function. We applied this analysis to characterize molecular mechanisms of pH-dependent allostery for the known pH sensor SHP2 (**Fig. 7A**). We find that physiological changes in pHi (0.1-0.4 pHi units) can produce 2-3 fold changes in basal SHP2 activity. We also used our computational pipeline to identify novel pH regulation of Src activity and identify the critical pH sensing residues (**Fig. 7B**). Importantly, we showed that the pH sensitive regulation of Src functions in concert with the well-characterized mode of phosphorylation (pY527) regulation. For example, simply lowering pHi can phenocopy Src activation growth factor stimulation and low pH can activate Src even when a phosphomimetic auto-inhibitory mutant is used (Src-Y527E). These data suggest that the transient pHi dynamics previously reported during cell cycle progression (0.1-0.2 pH units) (*2*, *46*), cell migration (∼0.3 pH units) (*4*, *47*, *48*) and cell differentiation (∼0.4 pH units) (*49*) may be sufficient to regulate basal cell signaling state of SHP2 and Src.

**Figure 7.**
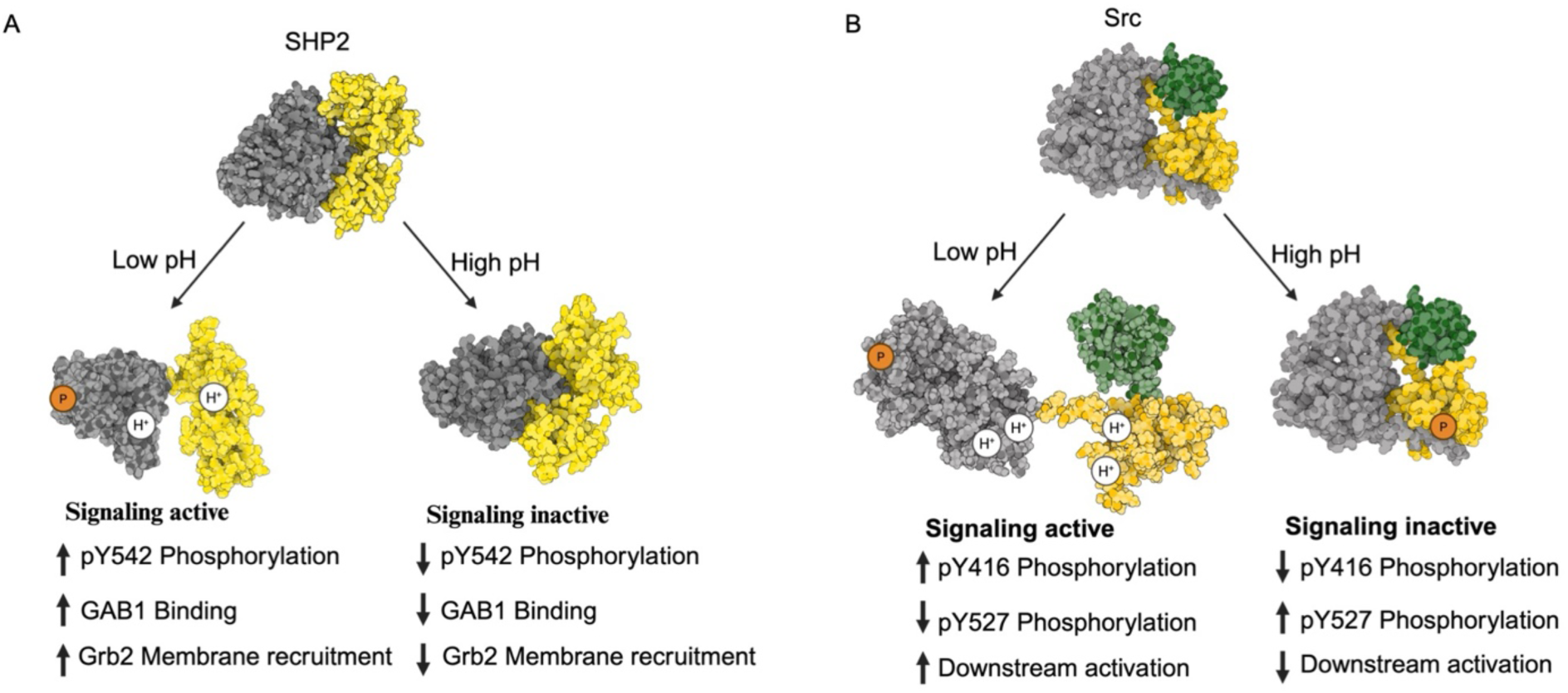
Molecular mechanism of pH-dependent allostery of SHP2 and Src. (A) Schematic of pH-driven activation and inhibition of SHP2. At low pH, the SH2 domain is unbound and SHP2 becomes signaling active with increased phosphorylation of Y542, increased GAB1 binding, and increased Grb2 recruitment. (B) Schematic of pH-driven activation and inhibition of Src. At low pH, the SH2 domain is unbound and Src becomes signaling active with increased phosphorylation of Y416, decreased phosphorylation of Y527, and increased substrate phosphorylation and downstream activation.

We also demonstrated that ionizable residues with predicted physiological pK_a_s are conserved across many SH2 domain-containing signaling proteins including kinases, phosphatases, transcription factors, and GTPase activating proteins. For multi-functional proteins that contain both evolutionarily conserved SH2 and SH3 domains, the residue networks with predicted physiological pKas clustered specifically at the interfaces between SH2 and the functional domains. This suggests that SH2 domains may be primed for pH sensitive allosteric regulation of signaling proteins. Our data showing the pH-dependence of Src under growth factor stimulation in cells and when using phosphomimetic mutants also suggests that pH-dependent regulation of SH2 signaling proteins may function in concert with traditional growth factor signaling and phosphorylation-dependent regulation. Future work will expand on this hypothesis with more mechanistic detail and functional validation.

We show that our prediction pipeline is as accurate as more computationally expensive prediction methods and does not require crystallographic waters or paths to solvent to identify residue networks. Our pipeline is also modular, enabling incorporation of improved pK_a_ prediction algorithms (e.g. recent machine learning (ML) pipelines with improved accuracy over PROPKA)(*50*). Most characterized pH-sensitive proteins have simple mechanisms where a single histidine residue titrates with changes in pH to confer changes in protein structure and function. We applied our network analysis to accurately identify previously validated pH-sensing residues based on existing structures, including in mutant p53 (His273)(*51*) and the sodium proton exchanger (H544)(*52*) (**Fig. S12**). We also showed that our pipeline accurately identified one of the only validated complex multi-site pH sensing mechanisms in isocitrate dehydrogenase 1 (D273, E304, K217)(*53*) (**Fig. S12**) this network was first reported using the more computationally expensive pHinder program to predict residues with physiological pKas (*53*).

Interestingly, recent work has suggested that His242 is required for pH-sensitive activity of phosphofructokinase muscle isoform (PFKM), but mutation of this residue did not completely abrogate the pH sensitive activity of PFKM (*54*). Our network analysis indicates that PFKM has two networked residues with physiological pKa (His242 and Asp309) (**Fig. S12**) and suggests that mutating both residues may fully abrogate pH-dependent PFKM function.

Scientists can apply our in silico approach to improve our mechanistic understanding of pH-dependent cell behaviors lacking known pH-sensitive molecular drivers, including differentiation(*49*, *55*), cell cycle progression(*2*, *3*), and epithelial to mesenchymal transition(*56*, *57*). In addition to revealing the pH-dependent structural proteome, this approach can be applied to molecularly characterize how dysregulated pHi drives pathophysiology in cancer (increased pHi)(*6*, *58*) and neurodegeneration (decreased pHi)(*59*).

## Materials and Methods

### pK_a_ prediction method

All full-length structures for proteins of interest were first identified based on the gene search and downloaded from the Protein Database (https://www.rcsb.org/). Next, the electrostatic properties for each structure were calculated using the APBS (Adaptive Poisson-Boltzmann Solver) webserver (https://server.poissonboltzmann.org/). The APBS calculation was performed using the following parameters: The pH for continuum electrostatic calculations was set to 7.4. The AMBER forcefield was used for the minimization step. We set the parameters in the APBS webserver to ensure that new atoms were not rebuilt too close to existing atoms and that hydrogen bonding networks were optimized during the structure repair and refinement step. The electrostatic data for each individual structure was combined into a single dataset and ionizable residues were then filtered for physiologically relevant pK_a_s. Histidine residues with pK_a_s above 7.0 were considered physiologically relevant while aspartates, glutamates, and lysines were filtered if pK_a_s were between 6.8.00. For ionizable residues that were classified using these criteria as “shifted” in ≥ 50% of structures, residue pK_a_s were averaged across all structures in the dataset. Next, the strength of coulombic interactions for the shifted residues were averaged for all structures. Structural visualization was done using Pymol and residue interaction networks were generated in Python (https://zenodo.org/records/15338058). For residue interaction networks, the length of the connecting lines indicates the strength of the coulombic interactions, with stronger coulombic interactions having shorter vectors.

### pK_a_ prediction benchmarking

A diverse set of experimental pK_a_ values were collected for Asp, Glu, His, Tyr and Lys from previous studies (*17*). The experimental dataset contained 34 proteins with 290 experimentally determined pK_a_s. Using our pipeline, we calculated the pK_a_s of these previously determined ionizable residues. We then calculated the weighted correlation coefficient for then predicted pK_a_s against the experimental pK_a_s and the Root Mean Squared Deviation (RMSD) of the predicted pK_a_ from the experimental values. The Garcia-Moreno lab have also calculated far-shifted pK_a_s for ionizable residues in Staphylococcal nuclease (SNase) (*18–20*). To determine if our pipeline was able to accurately predict these large pK_a_ shifts, we also generated a dataset comparing the delta pK_a_s of the validated dataset against our predictive set and calculated correlation coefficients and RMSDs from the experimental values.

### Structural conservation

The ConSurf server was used as a bioinformatics tool to estimate evolutionary conservation in proteins of interest. To determine the evolutionary conservation in our proteins of interest, we followed the recommended guidelines outlined by Yariv *et al.*, 2023(*33*). Briefly, the SHP2 (PDB: 2SHP) and Src (PDB:2SRC) protein structures were submitted to the ConSurf Web server (https://consurf.tau.ac.il/). ConSurf server utilizes an empirical Bayesian algorithm for conservation scores ranging from 1 to 9; where 1 is variable and 9 is conserved.

### Purification of SHP2 proteins

Full-length SHP2 variants were cloned into the pGEX-4TI SHP2 WT plasmid(*60*), pGEX-4T1 SHP2 WT was a gift from Ben Neel (Addgene plasmid # 8322; http://n2t.net/addgene:8322; RRID:Addgene_8322)) and expressed in *E. Coli* for purification. Transformed BL21(DE3) cells were grown in Miller’s Modified Luria Broth (RPI, L24080-2000) supplemented with 0.1 mg/mL ampicillin at 37°C with shaking (250 rpm). When cells reached an OD600 of 0.6, proteins were induced with Isopropyl β-d-1-thiogalactopyranoside (IPTG) (Thermo Scientific, R0392) (0.42 mM) and incubated at 18°C overnight with shaking (250 rpm). The induced cultures were centrifuged at 10,000 rpm at 4°C for 30 mins and resuspended in lysis buffer (B-PER™ Bacterial Protein Extraction Reagent (Thermo Scientific, 78243)) supplemented with EDTA-free Protease Inhibitor Cocktail (Roche, 04693132001)). Cell lysates were placed on a rocker at 4°C for 10 mins before centrifugating at 40,000 rpm at 4°C for 40 minutes. The supernatant was separated, and glutathione resin (GenScript, L00206) was added to bind the GST tagged proteins. The mixture was rocked for an hour at 4°C and then purified in a CrystalCruz chromatography column (Santa Cruz, sc-205554) at 4°C. Purified proteins were dialyzed into dialysis buffer (20mM Tris 50mM NaCl 1mM DTT 5% glycerol, pH 7.5) overnight at 4°C and stored at -20°C.

### Phosphatase activity assays

SHP2 variants were quantified by measuring absorbance at 280 nm and comparison with BSA protein standards. Para-nitrophenyl phosphate (pNPP) (Thermo Scientific, 34045) was used as an artificial substrate for the protein tyrosine phosphatase (PTPase) activity of each SHP2 variant.

Reactions were performed at 30°C in 100 μl volume of PTPase buffer (25 mM Tris–HCl, 50 mM NaCl, 2 mM EDTA and 10 mM dithiothreitol). Concentrations of pNPP ranged from 12-400 mM (12, 15.6, 25, 31.2, 50, 62.5, 100, 125, 250, 300, 400) and assay buffer pH values ranged from 6.1 to 8.0. All assay conditions were performed in technical duplicates in 96-well plates.

Reactions were initiated by the addition of 5μg of WT or mutant SHP2 to the reaction mixtures. Kinetic absorbance readings were collected at 405 nm over 30 mins on a spectrophotometer (Biotek Cytation 5). In all cases, after averaging technical replicates, the linear region of the reaction progress curve was determined by visual inspection and fit to a slope against the time of progression. These slopes were then plotted against the corresponding concentrations of substrate and fit to the Michaelis-Menten equation using non-linear regression to determine catalytic parameters. Experiments were repeated at least three times across two batches of purified proteins, and the average and standard error of the mean for all biological replicates are reported.

### Cell culture

Normal breast epithelial MCF10A cells were grown in complete media: DMEM/50% F12 w/ GlutaMax (Invitrogen, 10565-018) supplemented with 5% Horse Serum (Invitrogen, 16050-122), 0.02 µg/mL EGF (Peprotech, AF-100-15), 5 µg/mL Hydrocortisone (Sigma, H-0888), 10 mg/mL Insulin (Sigma, I-1882), 0.1 µg/mL Cholera toxin (Sigma, C-8052), and 1% Penicillin-Streptomycin (Corning, 30-001-Cl). Cells were maintained in humidified incubators at 37°C with 5% CO_2_. MCF10A cells (ATCC: CRL-10317) were split at sub-confluency. Cells were monitored for maintenance of epithelial morphology throughout experimental protocols. To lower pHi, complete media was supplemented with 25 μM EIPA (Invitrogen, E3111) and 30 μM S0859 (Sigma-Aldrich, SML0638-5MG). To raise pHi, complete media was supplemented with 30 mM ammonium chloride (NH_4_Cl, Fisher Chemical, A661-500, Lot 185503). To inhibit SHP2, complete media was supplemented with 15 μM PHPS1(Sigma-Aldrich, P0039) for 1 hour. To inhibit EGFR, complete media was supplemented with 1 μM Erlotinib (BioVision, 1588-1000) for 1 hour. To stimulate EGFR signaling, media was spiked with 10 ng/mL EGF (Preprotech, AF-100-15-100UG) for 5 mins. All chemical concentrations are final concentrations (Cf).

### pHi Manipulation and measurement

MCF10A cells were plated in 6 technical replicates for each treatment condition (6×100,000 cells/well; 24 well plate; 1 mL total volume). 24 hours after plating, media was exchanged with pHi manipulation media. After 1 hour of incubation with pHi manipulation media, pHi was measured as previously described(*61*). Briefly, 2′,7′-Bis-(2-Carboxyethyl)-5-(and-6)-Carboxyfluorescein, Acetoxymethyl (BCECF-AM) Ester (Biotium, 51011) was added at 2 μM (Cf) to cells for 30 min at 37°C with 5% CO_2_. Cells were then washed 3×5 min with HEPES-based wash buffer (30 mM HEPES, 145 mM NaCl, 5 mM KCl, 10 mM glucose, 1 mM KPO_4_, 1 mM MgSO_4_, and 2 mM CaCl_2_) at 37°C. Wash buffers were supplemented with appropriate pH-altering compounds at the same concentrations as the media. After washes, kinetic fluorescence measurements were made at pH-sensitive wavelengths (490ex/535em) and the pH-insensitive wavelengths (440ex/535em) every 15sec for 5 minutes at Medium PMT sensitivity on a plate reader (Biotek Cytation 5). After the initial kinetic pHi read, wash buffers were removed and replaced with the isotonic standardization buffer at ∼7.5 pH (25 mM HEPES, 80 mM KCl, 1 mM MgCl_2_) supplemented with the protonophore nigericin (Invitrogen, N1495) at 10 μM (Cf). Cells were incubated at 37°C for 10 min, and the plate was read again with the same parameters. The high pH standardization buffer was then replaced with a ∼7.0 pH nigericin buffer standard, cells were incubated at 37°C for 10 min, and the plate was read again with the same parameters. The nigericin buffer was then replaced with a ∼6.5 pH nigericin buffer standard, cells were incubated at 37°C for 10 min, and the plate was read again with the same parameters. Mean intensities from pHi measurement and the nigericin standards were exported and pHi was back calculated using a nigericin linear regression performed on each well with Nigericin standard pHi reported to the hundredths place.

### Western blotting

After pHi manipulation plates were removed from the incubator and placed on ice. The media was aspirated, and the plates were washed two times with ice-cold DPBS. 200µL of lysis buffer (500 mM NaCl, 10 mM EDTA, 500mM HEPES-free acid, 10 mM EGTA, 10% Triton X-100, protease inhibitor cocktail (Pierce™ Protease Inhibitor Tablets, A32965; 1 tablet/50 mL lysis buffer) and phosphatase inhibitor cocktail (PhosSTOP™, Sigma Aldrich, 4906845001), (pH 7.4) was added to each plate. After 15 minutes of incubation in the lysis buffer, cells were removed using cell scrapers and transferred to pre-chilled microfuge tubes. Lysates were centrifuged at 14,000 x g for 15 minutes at 4°C. Clarified lysates were isolated to separate microfuge tubes on ice. The Bradford assay was used to quantify the total protein content of lysates (Bradford, 1976) using Coomassie Protein reagent (Thermo Scientific, cat: 1856209). Lysates were stored at -80°C if not immediately being used for SDS-PAGE sample preparation.

Protein samples were prepared in 6X Laemilli (reducing) buffer (Thermo Scientific, J61337-AC) and boiled for 5 minutes at 100°C. A total of 10 µg of MCF10A lysate was added to each lane for immunoblot analysis. Protein samples were loaded onto 12% SDS-PAGE gels, which were run at 175V and 0.08 Amps for 15 minutes to stack samples. Protein samples were separated by running the gel at 200 V and 0.08 Amps for an hour and a half. Polyvinylidene difluoride (PVDF) membranes were pre-soaked in 100% methanol for 5 minutes and rinsed in turbo transfer buffer. Transfers were performed at 1.00 A and 25V for 30 mins using the TransBlot Turbo Transfer System (BioRad). Membranes were blocked for 1 hour in 5% milk (or 5% BSA for phospho-proteins) in 1X Tris-buffered Saline + 0.1% Tween® 20 (TBS-T) at room temperature with shaking. Membranes were washed 3×5 min in 1X TBS-T before being cut and incubated with primary antibodies. The following primary antibodies were used: SHP2 (Cell Signaling Technology, 3752; 1:1000 dilution in 1% milk in 1X TBS-T), pSHP2(Tyr524) (Cell Signaling Technology, 3751T; 1:1000 dilution in 1% BSA in 1X TBS-T), Cofilin (Cell Signaling Technology, 5175; 1:1000 dilution in 1% milk 1X TBS-T), HA (Cell Signaling Technology, 3724; 1:1000 dilution in 1% milk 1X TBS-T), Actin (Santa Cruz, sc58673; 1:1000 diluted in 1% milk in 1X TBS-T), Src (Cell Signaling Technology, 2109; 1:1000 dilution in 1% milk 1X TBS-T), pSrc(Tyr527) (Cell Signaling Technology, 2105; 1:1000 dilution in 1% BSA in 1X TBS-T), pSrc(Tyr416)( Cell Signaling Technology, 6943; 1:1000 dilution in 1% BSA in 1X TBS-T), ERK (Santa Cruz, sc-514302; 1:1000 diluted in 1% milk in 1X TBS-T), pERK (Cell Signaling Technology, 4370S; 1:1000 dilution in 1% BSA in 1X TBS-T), E-cadherin (Fisher Scientific, 13-190-0; 1:1000 diluted in 1% milk in 1X TBS-T), pE-cadherin (ThermoFisher Scientific, PA5-143661, 1:1000 diluted in 1% BSA in 1X TBS-T). Primary antibodies were incubated overnight at 4°C with shaking. Primary antibody solutions were removed, and membranes were washed 3×5 minutes in 1X TBS-T before incubating secondary antibodies (Goat anti-mouse/rabbit HRP-conjugated antibodies; 1:10,000 diluted in 1% milk/BSA in 1X TBS-T) for 1 hour at room temperature with shaking. Secondary antibody solutions were removed, and membranes were washed 3×5 minutes in 1X TBS-T before being developed using Pierce SuperSignal West Pico PLUS (Thermo Scientific, 34580) or Pierce SuperSignal West Femto PLUS (Thermo Scientific, 34095). Chemiluminescence and colorimetric images were acquired using a BioRad ChemiDoc imager.

Raw files were exported from ChemiDoc imager and opened in ImageJ. Band intensities were acquired by densitometry analysis (area under the curve of individual bands in respective lanes). For each replicate blot, target protein intensities were normalized to the intensity of loading controls before normalizing each condition ratio to that of the control within biological replicates.

### Co-immunoprecipitation assays

After pHi manipulation plates were washed 2x with ice-cold PBS. The cells were then lysed on ice. After Bradford measurements, 1µg of anti-SHP2 (Rabbit, Cell Signaling, 3752) was added to 100 µg of total lysate. The mixture was incubated on the rocker overnight at 4°C. Next, 80 µl of protein A-dynabeads were added to each tube and the mixture was incubated on the rocker for 4 hours at 4°C. Next the beads were washed 3 times with 800 µl of wash buffer (TBS + 0.1% Tween, EDTA-free Protease Inhibitor Cocktail (Roche, 04693132001)). After the final wash, the beads were heated at 50°C in 50 µL of 2x Laemilli sample buffer (Biorad, 1610737) for 5 minutes to elute the proteins. Next the immunoprecipitation samples and total lysates were immunoblotted. The following antibodies were used for primary incubations: SHP2(abcam, ab76285; 1:1000 dilution in 1% milk in 1X TBS-T), GAB1 (Santa Cruz Biotechnology, sc-133191, 1:1000 dilution in 1% milk in 1X TBS-T).

### GRB2 TagBFP Assay: Live-cell Microscopy

The pHR Grb2-TagBFP plasmid, a gift from Jared Toettcher (Addgene plasmid # 188631; http://n2t.net/addgene:188631; RRID:Addgene_188631)(*37*), was transfected into MCF10A cells and grown for 24 hours before plating for experiments. The cells were plated on a 35mm imaging dish with a 14mm glass coverslip (Matsunami, D35-14-1.5-U) a day before imaging. Cell membranes were visualized via CellMask Deep Red (Thermo Fisher, C10046; 1:20,000), incubated for 10 mins at 37°C in complete media. Microscope objectives were preheated to 37°C, and the stage-top incubator was preheated to 37°C and kept at 5% CO_2_/95% air. Confocal images were collected on a Nikon Ti-2 spinning disk confocal with a 60× (CFI PLAN FLUOR NA 1.3) oil immersion objective. The microscope is equipped with a stage-top incubator (Tokai Hit), a Yokogawa spinning disk confocal head (CSU-X1), four laser lines (405 nm, 488 nm, 561 nm and 647 nm), appropriate filter sets (BFP: 455/50; GFP: 525/36; Tx red: 605/52, and Cy5: 705/72), a Ti2-S-SE motorized stage, multi-point perfect focus system and an Orca Flash 4.0 CMOS camera. The cells were imaged initially for starting intensities before nigericin stimulation. Next, the media was replaced with nigericin buffer (see **pHi manipulation and measurement** section) to pin the cells at different pH values. The BFP (405 ex, 455/50 em; 500ms exposure) and Cell Mask (647ex, 705/72 em; 300ms exposure) for 15 mins at 30 second intervals. The image quantification was performed in Nikon Elements software. Images were background-subtracted using a region of interest (ROI) drawn on a glass coverslip (determined by DIC). To only measure the intensity of the cell membrane, the ROI for membrane of each cell was outlined using the membrane marker and tracked over time. In case of improper tracking over time points, manual tracking was used to redraw ROIs.

### Immunofluorescence assays

MCF10A Cells were plated in 35mm imaging dish with a 14mm glass coverslip (Matsunami, D35-14-1.5-U) a day before treatment. The cells were then treated with pHi manipulation drugs for an hour. Cells were rinsed with DPBS and fixed in 3.7% formaldehyde (Alfa Aestar, 33314) at room temperature for 10 min. Cells were washed three times for 2 min each time with DPBS, then incubated in lysis buffer (0.1% Triton-X in DPBS) for 10 min at room temperature. Cells were washed three times for 2 min each time with DPBS, then incubated in blocking buffer (1.0% BSA in DPBS) for 1 hour at room temperature with rocking followed by three 2 min washes in DPBS. Cells were then incubated overnight at 4°C with anti-C-Src antibody (Proteintech, 60315-1-Ig) and anti-pY416-C-Src antibody (ThermoFisher Scientific, 44-660G). Cells were washed three times for 3 min each time in DPBS, then incubated with secondary antibody and phalloidin for 1 hour at room temperature. After three 3 min washes in DPBS, Hoechst 33342 dye (1:20,000) in antibody buffer was added for 10 min, then removed. Cells were imaged in DPBS on the spinning disk confocal microscope. Images were collected in BFP (405 nm laser excitation, 455/50 nm emission, 200-500msec exposures); GFP (488 nm laser excitation, 525/36 nm emission, 200-500msec exposures); Tx red (561 nm laser excitation, 605/52 nm emission, 200-500msec exposures); and Cy5(647 nm laser excitation, 705/72 nm emission, 200-500msec exposures).

### Constant pH molecular dynamics

The 2SHP and 2SRC monomer chains were downloaded from Protein Data Bank (PDB) for initial structures. Constant pH MD (CpHMD) simulations were conducted following the protocol described by Henderson *et al*., 2022(*34*). CpHMD simulations were performed with implicit solvent using the GBNeck2-CpHMD method implemented in AMBER 2022(*62*). Preparation of input files was done using the cphmd_pred.sh script provided in the protocol(*34*), using the default titratable residue types (Asp, Glu, His, Cys, Lys). After energy minimization, structure relaxation, and protonation state initiation(*34*), independent production simulations were run over 4.0-10.0 pH in half-integer increments for 8 nanoseconds (ns) for each, resulting in 13 independent CpHMD simulations and 104 ns of total simulation time for each structure.

Coordinate snapshots were saved every 20 picoseconds (ps), and protonation state snapshots every 2 ps. The probability of protonated states became time-independent after 4 ns. The resulting coordinate and protonation state trajectories were analyzed using Python scripts provided in the protocol(*62*), the CPPTRAJ analysis program(*63*), and in-house Python scripts (https://zenodo.org/records/15338058). To create the initial structure for the H116AE252A mutant, we introduced the H116A and E252A mutations into the wild-type PDB (2SHP) using the Mutagenesis Wizard functionality in Pymol Version 2.6. This mutant was simulated using the same CpHMD protocol as for the WT protein.

### Statistical analysis

GraphPad Prism was used to prepare graphs and perform statistical analyses. Normality tests were performed on all data sets as well as outlier tests using the ROUT method (Q=1%).

Normally distributed data is shown with means and non-normally distributed data is shown with medians. For normally distributed data, significance was determined by ratio paired t-tests or appropriate ANOVA. For non-normally distributed data, significance was determined by Kruskal-Wallis test. In all statistical tests, appropriate multiple comparisons corrections were performed. “N” indicates the number of biological replicates performed and “n” represents the number of technical replicates or individual cell measurements collected.

## Supporting information

Video1_2shp_pH4.0

Video2_2shp_pH4.5

Video3_2shp_pH5.0

Video4_2shp_pH5.5

Video5_2shp_pH6.0

Video6_2shp_pH6.5

Video7_2shp_pH7.0

Video8_2shp_pH7.5

Video9_2shp_pH8.0

Video10_2shp_pH8.5

Video11_2shp_pH9.0

Video12_2shp_pH9.5

Video13_2shp_pH10.0

Video14_2src_pH4.0

Video15_2src_pH4.5

Video16_2src_pH5.0

Video17_2src_pH5.5

Video18_2src_pH6.0

Video19_2src_pH6.5

Video20_2src_pH7.0

Video21_2src_pH7.5

Video22_2src_pH8.0

Video23_2src_pH8.5

Video24_2src_pH9.0

Video25_2src_pH9.5

Video26_2src_pH10.0

## Acknowledgements

This research was supported in part by the Notre Dame Center for Research Computing; we specifically thank Dr. Dodi Heryadi and Dr. Scott Hampton for technical support.

## Author contributions

PKVD contributed to project conceptualization, methodology development, data collection, analysis, figure generation, writing (original draft and review/editing).

LP contributed to methodology development, data collection, analysis, figure generation, and writing (original draft and review/editing.

EAG contributed to data collection, analysis, figure generation, writing (review/editing). ETN contributed to data collection, analysis, figure generation, writing (review/editing). JAA contributed to data collection, analysis, writing (review/editing).

LMH contributed to contributed to data collection, analysis, writing (review/editing). JWP contributed to writing (original draft and review/editing), supervision, and funding acquisition.

KAW contributed to project conceptualization, methodology development, analysis, figure generation, writing (original draft and review/editing), project administration, supervision, and funding acquisition.

## Materials & Correspondence

For materials requests and correspondence contact Katharine White

## References

1. Boron, W. F. Regulation of intracellular pH. Adv. Physiol. Educ. 28, 160–179 (2004).

2. Spear, J. S. & White, K. A. Single-cell intracellular pH dynamics regulate the cell cycle by timing the G1 exit and G2 transition. J. Cell Sci. 136, jcs260458 (2023).

3. Putney, L. K. & Barber, D. L. Na-H Exchange-dependent Increase in Intracellular pH Times G2/M Entry and Transition. J. Biol. Chem. 278, 44645–44649 (2003).

4. Frantz, C., Karydis, A., Nalbant, P., Hahn, K. M. & Barber, D. L. Positive feedback between Cdc42 activity and H+ efflux by the Na-H exchanger NHE1 for polarity of migrating cells. J. Cell Biol. 179, 403–410 (2007).

5. Boussouf, A. & Gaillard, S. Intracellular pH changes during oligodendrocyte differentiation in primary culture. J. Neurosci. Res. 59, 731–739 (2000).

6. Czowski, B. J., Romero-Moreno, R., Trull, K. J. & White, K. A. Cancer and pH Dynamics: Transcriptional Regulation, Proteostasis, and the Need for New Molecular Tools. Cancers 12, 2760 (2020).

7. Garcia-Moreno, B. Adaptations of proteins to cellular and subcellular pH. J. Biol. 8, 98 (2009).

8. Harms, M. J. et al. A buried lysine that titrates with a normal pKa: role of conformational flexibility at the protein-water interface as a determinant of pKa values. Protein Sci 17, 833– 45 (2008).

9. Harms, M. J. et al. The pK(a) values of acidic and basic residues buried at the same internal location in a protein are governed by different factors. J Mol Biol 389, 34–47 (2009).

10. Isom, D. G., Castaneda, C. A., Cannon, B. R. & Garcia-Moreno, B. Large shifts in pKa values of lysine residues buried inside a protein. Proc Natl Acad Sci U A 108, 5260–5 (2011).

11. Isom, D. G. & Dohlman, H. G. Buried ionizable networks are an ancient hallmark of G protein-coupled receptor activation. Proc. Natl. Acad. Sci. 112, 5702–5707 (2015).

12. Isom, D. G. et al. Protons as Second Messenger Regulators of G Protein Signaling. Mol. Cell 51, 531–538 (2013).

13. Tews, I. et al. The Structure of a pH-Sensing Mycobacterial Adenylyl Cyclase Holoenzyme. Science 308, 1020–1023 (2005).

14. Olsson, M. H. M., Søndergaard, C. R., Rostkowski, M. & Jensen, J. H. PROPKA3: Consistent Treatment of Internal and Surface Residues in Empirical p *K* _a_ Predictions. J. Chem. Theory Comput. 7, 525–537 (2011).

15. Bertalan, É., Rodrigues, M. J., Schertler, G. F. X. & Bondar, A. Graph-based algorithms to dissect long-distance water-mediated H-bond networks for conformational couplings in GPCRs. Br. J. Pharmacol. bph.16387 (2024) doi:10.1111/bph.16387.

16. Qu, C. K. The SHP-2 tyrosine phosphatase: Signaling mechanisms and biological functions. Cell Res. 10, 279–288 (2000).

17. Mannell, H. & Krotz, F. SHP-2 Regulates Growth Factor Dependent Vascular Signalling and Function. Mini-Rev. Med. Chem. 14, 471–483 (2014).

18. Lauriol, J., Jaffré, F. & Kontaridis, M. I. The role of the protein tyrosine phosphatase SHP2 in cardiac development and disease. Semin. Cell Dev. Biol. 37, 73–81 (2015).

19. Merritt, R., Hayman, M. J. & Agazie, Y. M. Mutation of Thr466 in SHP2 abolishes its phosphatase activity, but provides a new substrate-trapping mutant. Biochim. Biophys. Acta BBA - Mol. Cell Res. 1763, 45–56 (2006).

20. Schaffhausen, B. SH2 domain structure and function. Biochim. Biophys. Acta BBA - Rev. Cancer 1242, 61–75 (1995).

21. Hof, P., Pluskey, S., Dhe-Paganon, S., Eck, M. J. & Shoelson, S. E. Crystal Structure of the Tyrosine Phosphatase SHP-2. Cell 92, 441–450 (1998).

22. Srivastava, J. et al. Structural model and functional significance of pH-dependent talin-actin binding for focal adhesion remodeling. Proc Natl Acad Sci U A 105, 14436–41 (2008).

23. Vercoulen, Y. et al. A histidine pH sensor regulates activation of the Ras-specific guanine nucleotide exchange factor RasGRP1. Elife 6, e29002 (2017).

24. Yariv, B. et al. Using evolutionary data to make sense of macromolecules with a “face-lifted” ConSurf. Protein Sci. 32, e4582 (2023).

25. Henderson, J. A. et al. A Guide to the Continuous Constant pH Molecular Dynamics Methods in Amber and CHARMM [Article v1.0]. Living J. Comput. Mol. Sci. 4, (2022).

26. Sun, J. et al. Antagonism between binding site affinity and conformational dynamics tunes alternative cis-interactions within Shp2. Nat. Commun. 4, 2037 (2013).

27. Cai, T., Nishida, K., Hirano, T. & Khavari, P. A. Gab1 and SHP-2 promote Ras/MAPK regulation of epidermal growth and differentiation. J. Cell Biol. 159, 103–112 (2002).

28. Farahani, P. E. et al. pYtags enable spatiotemporal measurements of receptor tyrosine kinase signaling in living cells. eLife 12, e82863 (2023).

29. Hellmuth, K. et al. Specific inhibitors of the protein tyrosine phosphatase Shp2 identified by high-throughput docking. Proc. Natl. Acad. Sci. 105, 7275–7280 (2008).

30. Singer, A. U. & Forman-Kay, J. D. pH Titration studies of an SH2 domain-phosphopeptide complex: Unusual histidine and phosphate p *K _a_* values. Protein Sci. 6, 1910–1919 (1997).

31. 31. Vogel, H. J. [13] Phosphorus-31 nuclear magnetic resonance of phosphoproteins. in Methods in Enzymology vol. 177 263–282 (Elsevier, 1989).

32. Kurochkina, N. & Guha, U. SH3 domains: modules of protein–protein interactions. Biophys. Rev. 5, 29–39 (2013).

33. Xu, W., Doshi, A., Lei, M., Eck, M. J. & Harrison, S. C. Crystal Structures of c-Src Reveal Features of Its Autoinhibitory Mechanism. Mol. Cell 3, 629–638 (1999).

34. Sondka, Z. et al. COSMIC: a curated database of somatic variants and clinical data for cancer. Nucleic Acids Res. 52, D1210–D1217 (2024).

35. White, K. A. et al. Cancer-associated arginine-to-histidine mutations confer a gain in pH sensing to mutant proteins. Sci. Signal. 10, eaam9931 (2017).

36. Kisor, K. pH-Sensors Regulating Transcription, Metabolism, and Cancer Cell Biology. (2023).

37. Luna, L. A. et al. An acidic residue buried in the dimer interface of isocitrate dehydrogenase 1 (IDH1) helps regulate catalysis and pH sensitivity. Biochem. J. 477, 2999–3018 (2020).

38. Webb, B. A. et al. A Histidine Cluster in the Cytoplasmic Domain of the Na-H Exchanger NHE1 Confers pH-sensitive Phospholipid Binding and Regulates Transporter Activity. J Biol Chem 291, 24096–24104 (2016).

39. Chen, A. Y., Lee, J., Damjanovic, A. & Brooks, B. R. Protein p *K* _a_ Prediction by Tree-Based Machine Learning. J. Chem. Theory Comput. 18, 2673–2686 (2022).

40. Ulmschneider, B. et al. Increased intracellular pH is necessary for adult epithelial and embryonic stem cell differentiation. J Cell Biol 215, 345–355 (2016).

41. Liu, Y. et al. Intracellular pH dynamics regulates intestinal stem cell lineage specification. Nat. Commun. 14, 3745 (2023).

42. Amith, S. R., Wilkinson, J. M. & Fliegel, L. Na+/H+ exchanger NHE1 regulation modulates metastatic potential and epithelial-mesenchymal transition of triple-negative breast cancer cells. Oncotarget 7, 21091–113 (2016).

43. Suzuki, A., Maeda, T., Baba, Y., Shimamura, K. & Kato, Y. Acidic extracellular pH promotes epithelial mesenchymal transition in Lewis lung carcinoma model. Cancer Cell Int. 14, (2014).

44. White, K. A., Grillo-Hill, B. K. & Barber, D. L. Cancer cell behaviors mediated by dysregulated pH dynamics at a glance. J. Cell Sci. 130, 663–669 (2017).

45. Harguindey, S., Reshkin, S. J., Orive, G., Arranz, J. L. & Anitua, E. Growth and trophic factors, pH and the Na+/H+ exchanger in Alzheimer’s disease, other neurodegenerative diseases and cancer: new therapeutic possibilities and potential dangers. Curr Alzheimer Res 4, 53–65 (2007).

46. O’Reilly, A. M., Pluskey, S., Shoelson, S. E. & Neel, B. G. Activated Mutants of SHP-2 Preferentially Induce Elongation of *Xenopus* Animal Caps. Mol. Cell. Biol. 20, 299–311 (2000).

47. Choi, C.-H., Webb, B. A., Chimenti, M. S., Jacobson, M. P. & Barber, D. L. pH sensing by FAK-His58 regulates focal adhesion remodeling. J. Cell Biol. 202, 849–859 (2013).

48. Case, D. A. et al. Amber 2022. Unpublished 10.13140/RG.2.2.31337.77924 (2022).

49. Roe, D. R. & Cheatham, T. E. PTRAJ and CPPTRAJ: Software for Processing and Analysis of Molecular Dynamics Trajectory Data. J. Chem. Theory Comput. 9, 3084–3095 (2013).

