## Supplementary Figures for "Ionizable networks mediate pH-dependent allostery in SH2 signaling proteins"

A

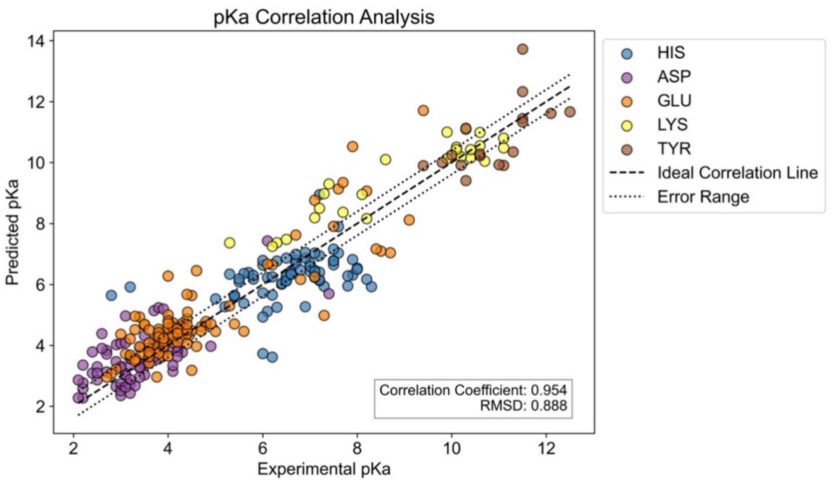


B

| Res | Correlation Coefficient | RMSD |
|---|---|---|
| HIS | 0.302 | 1.025 |
| ASP | 0.816 | 0.714 |
| GLU | 0.864 | 0.891 |
| LYS | 0.927 | 0.950 |
| TYR | 0.746 | 0.848 |

C

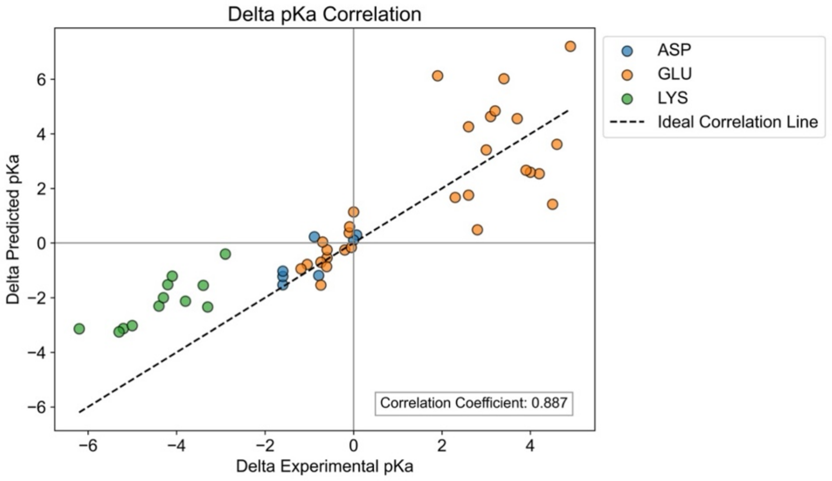


D

| Res | RMSD |
|---|---|
| Total | 1.632 |
| ASP | 0.527 |
| GLU | 2.243 |
| LYS | 1.503 |

**Fig. S1. Benchmarking of in silico pKa prediction platform.** (A) Predicted pKa values for experimentally calculated values were determined for 34 well-characterized pH sensors with 290 known pKas. We generated correlation plots for predicted pKa values against the experimentally calculated pKa values. The Correlation coefficient for our prediction was 0.954. The RMSD was 0.888 pH units. (B) Table of the correlation coefficient and RMSD for His, Asp, Glu, Lys, and Tyr residues in the correlation plot in (A). (C) Correlation between predicted and experimental pKa values for SNase mutants with large pKa shifts. The correlation coefficient was 0.887. (D) Table of RMSD values for Total, Asp, Glu, Lys ionizable residues in the correlation plot in (C).

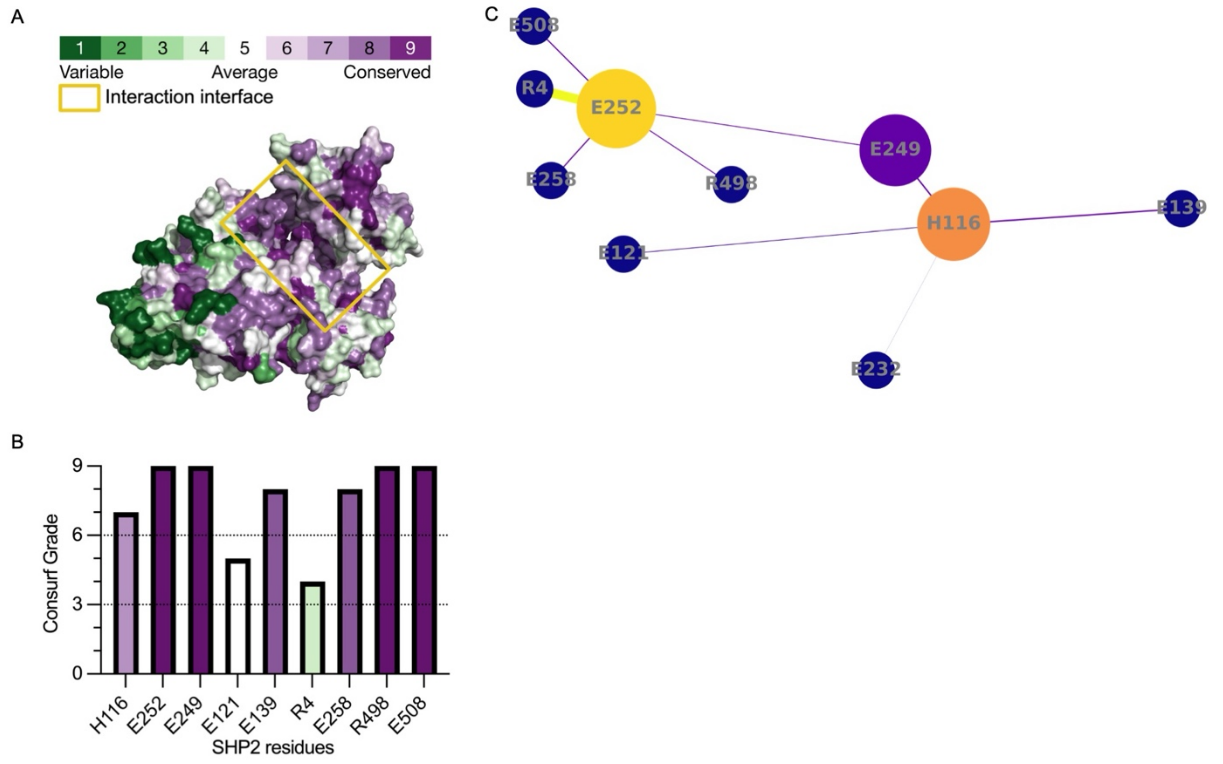


**Fig. S2. Identified residue network in SHP2 is evolutionarily and structurally conserved.** (A) Consurf surface structure of SHP2 (2SHP) with conservation scores for each amino acid from variable to conserved. (B) Consurf grades for shifted residues and their coulombic interactors found in the predicted network. (C) Residue interaction network of shifted ionizable residues and their coulombic interactors. The color spectrum ranges from dark purple to bright yellow. Nodes become more yellow as their degree increases. Larger nodes indicate higher betweenness centrality (measure of how often a node lies on the shortest path between all pairs of nodes in a network). Edges that are yellower and wider represent a higher interaction score.

A

| WT | $K_m$ (M) | $K_{cat}$ ($s^{-1}$) | Cat Efficiency ($M^{-1}s^{-1}$) |
|---|---|---|---|
| 6.1 | 0.09±0.01 | 0.34±0.01 | 3.97±0.45 |
| 6.4 | 0.10±0.01 | 0.64±0.02 | 6.71±0.65 |
| 6.7 | 0.11±0.01 | 0.87±0.04 | 8.04±1.04 |
| 7 | 0.10±0.01 | 0.69±0.04 | 6.74±0.98 |
| 7.2 | 0.09±0.02 | 0.61±0.06 | 6.64±1.87 |
| 7.4 | 0.09±0.01 | 0.54±0.03 | 5.72±0.86 |
| 7.7 | 0.10±0.02 | 0.42±0.03 | 4.09±0.74 |
| 8 | 0.09±0.01 | 0.37±0.01 | 4.07±0.41 |
| **E252** | | | |
| 6.1 | 0.11±0.01 | 0.85±0.03 | 7.66±0.70 |
| 6.4 | 0.11±0.02 | 0.86±0.06 | 7.77±1.36 |
| 6.7 | 0.12±0.02 | 0.81±0.05 | 6.62±1.01 |
| 7 | 0.11±0.01 | 0.49±0.02 | 4.70±0.57 |
| 7.2 | 0.12±0.01 | 0.49±0.02 | 4.05±0.39 |
| 7.4 | 0.12±0.02 | 0.52±0.03 | 4.35±0.62 |
| 7.7 | 0.12±0.03 | 0.54±0.06 | 4.55±1.36 |
| 8 | 0.11±0.03 | 0.44±0.05 | 3.88±1.14 |
| **H116** | | | |
| 6.1 | 0.10±0.01 | 0.80±0.02 | 7.99±0.61 |
| 6.4 | 0.10±0.02 | 0.81±0.05 | 7.92±1.27 |
| 6.7 | 0.11±0.02 | 0.81±0.06 | 7.26±1.44 |
| 7 | 0.12±0.02 | 0.80±0.06 | 6.50±1.29 |
| 7.2 | 0.13±0.01 | 0.73±0.02 | 5.59±0.46 |
| 7.4 | 0.11±0.02 | 0.67±0.04 | 6.00±0.93 |
| 7.7 | 0.08±0.01 | 0.43±0.03 | 5.62±1.15 |
| 8 | 0.09±0.02 | 0.40±0.03 | 4.71±1.11 |
| **E249** | | | |
| 6.1 | 0.11±0.03 | 0.50±0.05 | 4.44±1.13 |
| 6.4 | 0.13±0.02 | 0.60±0.04 | 4.64±0.85 |
| 6.7 | 0.10±0.03 | 0.79±0.08 | 7.62±2.08 |
| 7 | 0.09±0.02 | 0.78±0.06 | 8.62±1.82 |
| 7.2 | 0.09±0.01 | 0.66±0.03 | 7.39±1.03 |
| 7.4 | 0.07±0.01 | 0.60±0.03 | 8.40±1.06 |
| 7.7 | 0.09±0.01 | 0.53±0.02 | 6.02±0.75 |
| 8 | 0.07±0.01 | 0.33±0.02 | 4.66±0.88 |
| **H116A/E252A** | | | |
| 6.1 | 0.10±0.01 | 0.62±0.03 | 6.28±0.74 |
| 6.4 | 0.10±0.01 | 0.58±0.03 | 6.01±0.75 |
| 6.7 | 0.10±0.02 | 0.60±0.04 | 6.13±1.06 |
| 7 | 0.11±0.02 | 0.65±0.04 | 5.68±0.96 |
| 7.2 | 0.09±0.01 | 0.56±0.03 | 6.09±0.78 |
| 7.4 | 0.11±0.01 | 0.63±0.02 | 5.76±0.39 |
| 7.7 | 0.11±0.02 | 0.61±0.04 | 5.35±0.99 |
| 8 | 0.12±0.03 | 0.60±0.06 | 5.05±1.24 |

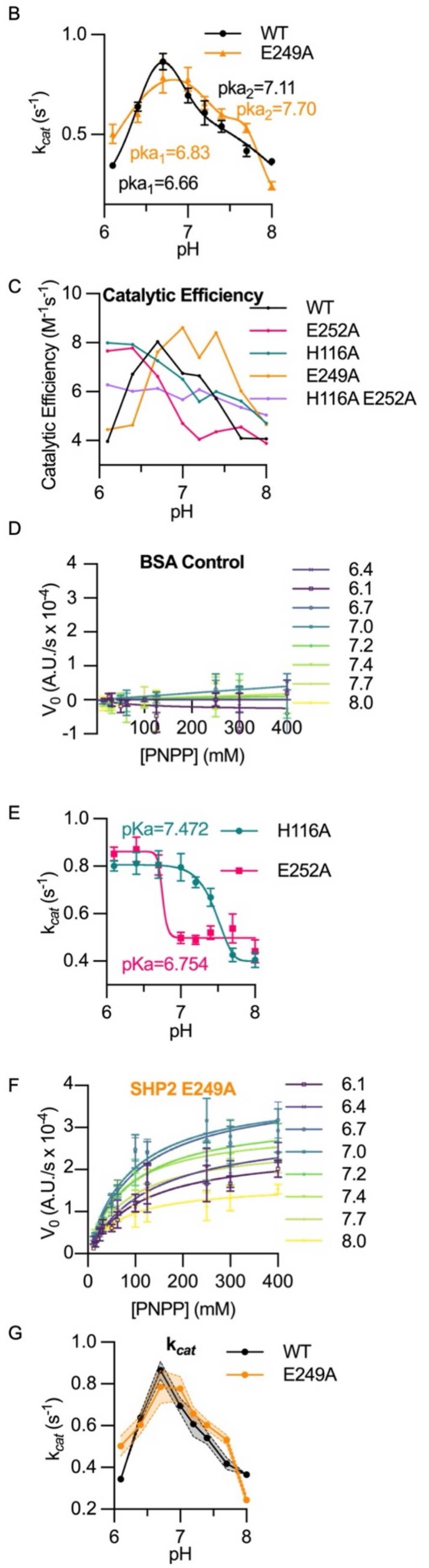


**Fig. S3. Enzyme kinetics collected from SHP2 wildtype and mutants.** (A) Calculated $K_m$, $k_{cat}$, and Catalytic Efficiency ($k_{cat}$/ $K_m$) values for pH conditions 6.1-8.0 for WT, E252A, H116A, E249A, and H116A E252A SHP2. (mean ± SEM, N=3) calculated from data collected as in Fig. 2. (B) Plot of $k_{cat}$ vs. pH for WT and E249A SHP2 activity and fit to a sum of two gaussians to determine apparent $pK_a$ values. (C) Plot of catalytic efficiency vs. pH for WT, H116A, E252A, E249A, and H116A E252A SHP2. (mean ± SEM, N=3) calculated from data collected as in Fig. 2. (D) Control experiment using BSA in vitro phosphatase activity, assays performed as in Fig. 2A. (mean ± SEM, N=3). (E) Apparent $pK_a$ fits of H116A and E252A SHP2 calculated from plot of $k_{cat}$ vs. pH for H116A and E252A SHP2 activity. (mean ± SEM, N=3) (F) Single-mutant E249A SHP2 in vitro phosphatase activity, assays performed as described for Fig. 2A. (mean ± SEM; N=3 from ≥2 different protein preparations). (G) Plot of $k_{cat}$ vs. pH for WT and E249A SHP2 activity. Calculated from activity curves in A, D and E. (mean ± SEM, N=3).

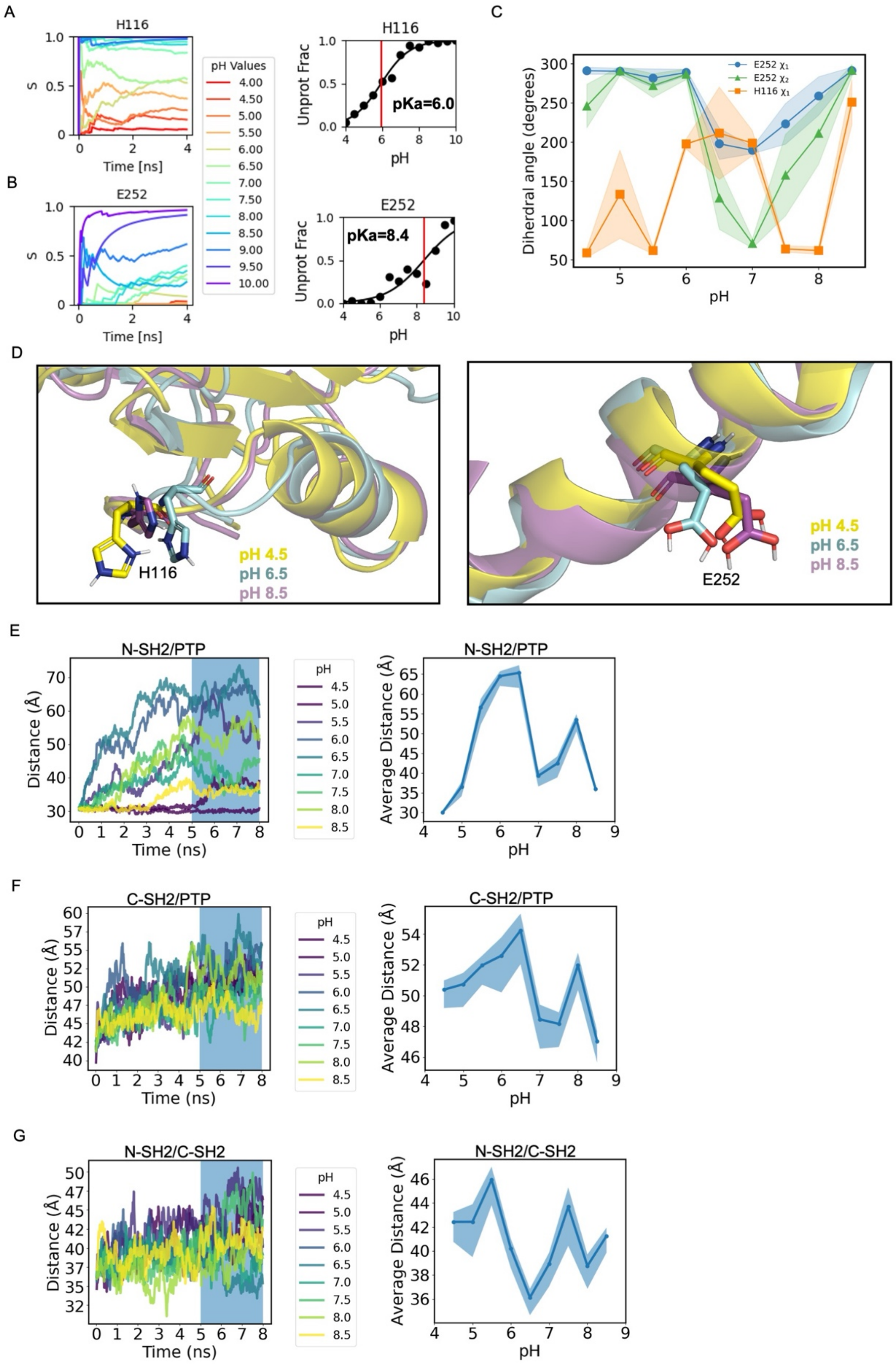

A
H116
S
Time [ns]
pH Values
4.00
4.50
5.00
5.50
6.00
6.50
7.00
7.50
8.00
8.50
9.00
9.50
10.00
H116
Unprot Frac
pKa=6.0
pH
B
E252
S
Time [ns]
E252
Unprot Frac
pKa=8.4
pH
C
Diherdral angle (degrees)
E252 $\chi_1$
E252 $\chi_2$
H116 $\chi_1$
pH
D
H116
pH 4.5
pH 6.5
pH 8.5
E252
pH 4.5
pH 6.5
pH 8.5
E
N-SH2/PTP
Distance (Å)
Time (ns)
pH
4.5
5.0
5.5
6.0
6.5
7.0
7.5
8.0
8.5
N-SH2/PTP
Average Distance (Å)
pH
F
C-SH2/PTP
Distance (Å)
Time (ns)
pH
4.5
5.0
5.5
6.0
6.5
7.0
7.5
8.0
8.5
C-SH2/PTP
Average Distance (Å)
pH
G
N-SH2/C-SH2
Distance (Å)
Time (ns)
pH
4.5
5.0
5.5
6.0
6.5
7.0
7.5
8.0
8.5
N-SH2/C-SH2
Average Distance (Å)
pH

**Fig. S4. Constant pH molecular dynamics (CpHMD) reveals pH-dependent release of SH2 domains in WT SHP2.** (A and B) Time series from CpHMD showing protonation states of indicated residues for the first 4 ns for pH values 4.0 to 10.0 (left) and titration plot and $pK_a$ (right) for H116 (A) and E542 (B). (C) Plot of dihedral angles of H116 and E252 for the last 3 ns for pH values 4.5 to 8.5. (D) Pair-fit views of zoomed-in structures of SHP2 showing the stick views of H116 and E252 at 8 ns for pH 4.5 (yellow), 6.5 (cyan), and 8.5 (magenta). (E to G) Time series from CpHMD showing interdomain distances for pH values 4.5-8.5 (left) and average interdomain distance (over the last 3 ns of simulation time) (right) (mean ± SD) for (E) N-terminal SH2 (N-SH2) and protein tyrosine phosphatase (PTP), (F) C-terminal SH2 (C-SH2) and PTP domains and (G) N-SH2 and C-SH2 domains of SHP2.

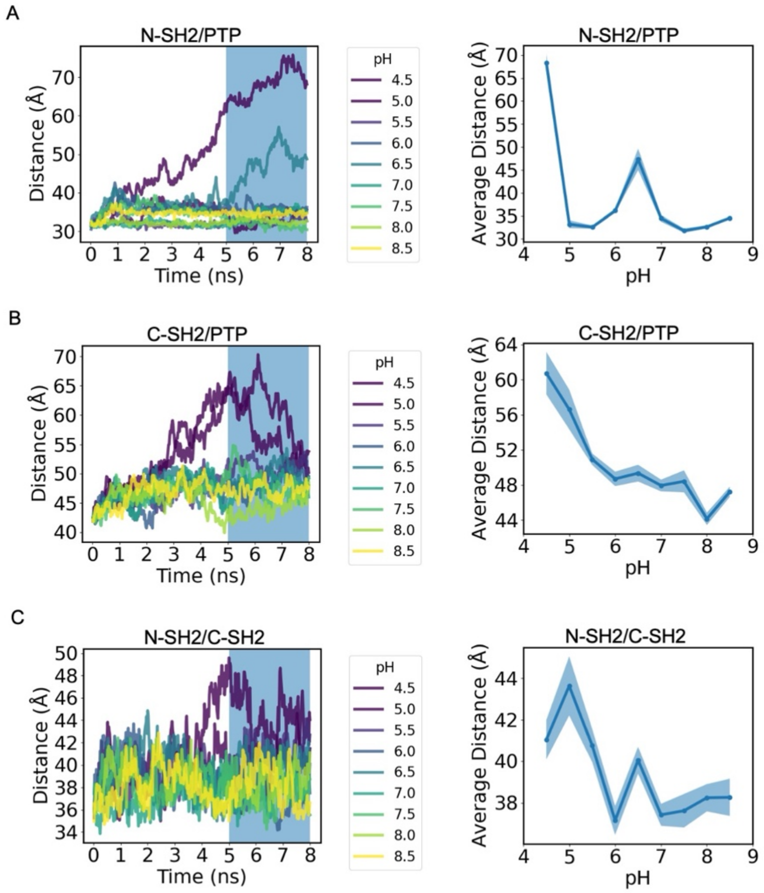


**Fig. S5. Constant pH molecular dynamics (CpHMD) reveals pH-dependent release of SH2 domains in H116A E252A SHP2 is abrogated** (A to C) Time series from CpHMD showing interdomain distances for pH values 4.5-8.5 (left) and average interdomain distance (over the last 3 ns of simulation time) (right) (mean ± SD) for (A) N-terminal SH2 (N-SH2) and protein tyrosine phosphatase (PTP), (B) C-terminal SH2 (C-SH2) and PTP domains and (C) N-SH2 and C-SH2 domains of SHP2.

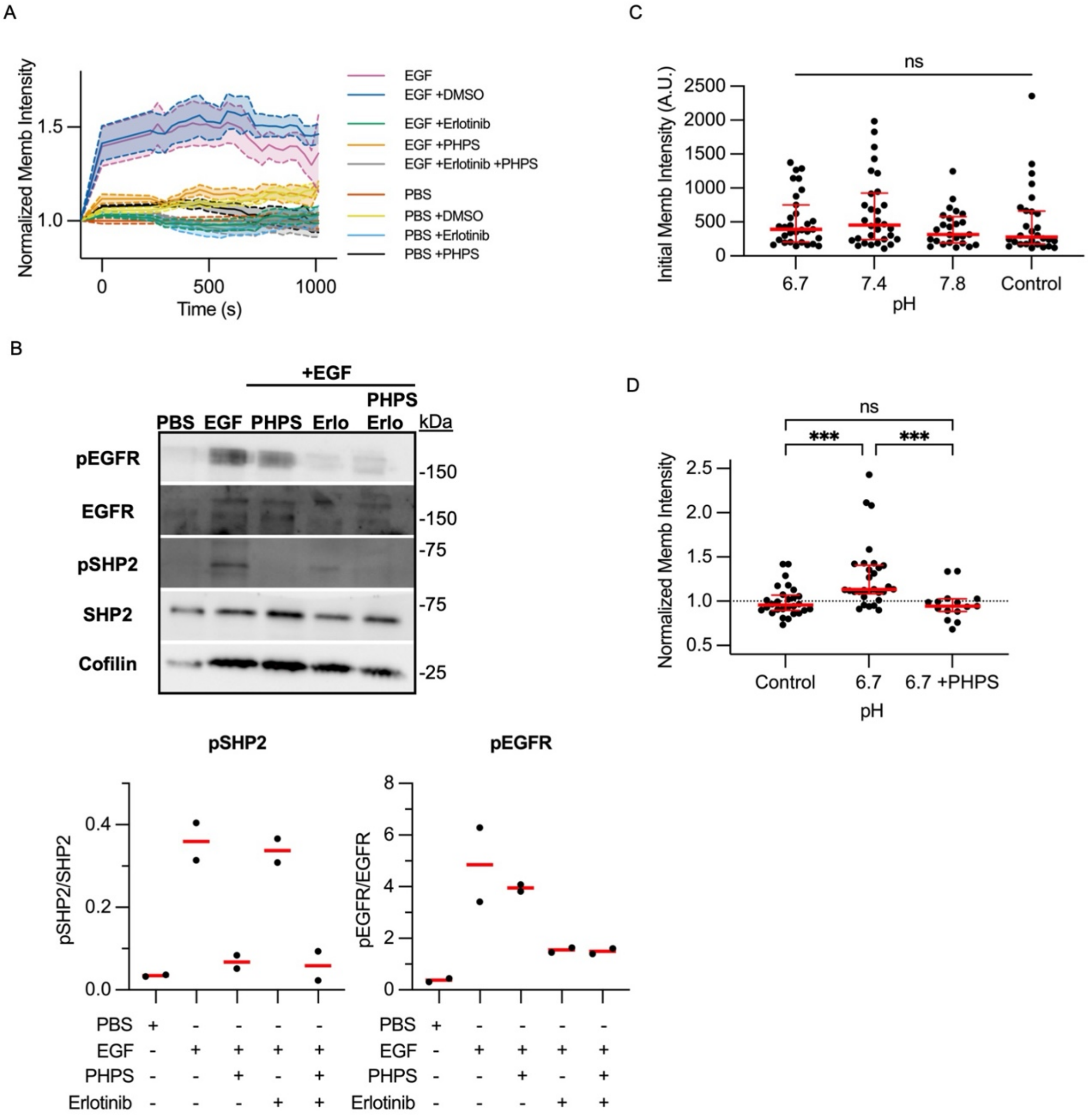


**Fig. S6. Grb2-TagBFP is a specific biosensor for SHP2 activity.** (A) Quantification of membrane intensity of Grb2 TagBFP with and without SHP2 inhibitor (PHPS), EGFR inhibitor (Erlotinib), and controls (DMSO and PBS) with and without EGF stimulation. Intensities were photobleach-corrected and normalized to PBS control. (mean ± SEM, N=3) (B) Representative immunoblot of pEGFR (pY1068) and pSHP2 (pY542) for EGF/PBS stimulated serum-starved MCF10A cells when treated with PHPS, Erlotinib, both PHPS and Erlotinib or control. Quantification of replicate experiments for pSHP2 (pY542) on left and pEGFR (pY1068) on right. (N=2). (C) Quantification of initial membrane intensities of single cells described in Fig 3e-g. (6.7 pH, n=30, 7.4 pH, n=30, 7.8 pH, n=25, control, n=28). (D) End point quantification of membrane intensities of single cells stimmed at pH of 6.7 with or without SHP2 inhibitor (PHPS). For c and d, significance was determined using the Kruskal-Wallis test. * $p<0.05$, ** $p<0.01$, *** $p<0.001$, **** $p<0.0001$.

A

**PTN6 (SHP1)**

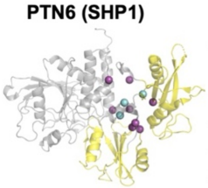

| Residue | Predicted $pK_a$ |
|---|---|
| Glu247 | 7.1 ± 0.0 |
| His114 | 7.3 ± 0.0 |
| Glu249 | 8.9 ± 0.0 |

B

**ZAP70**

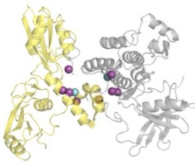

| Residue | Predicted $pK_a$ |
|---|---|
| Glu150 | 6.3 ± 0.1 |
| Glu440 | 7.8 ± 1.7 |

C

**JAK1**

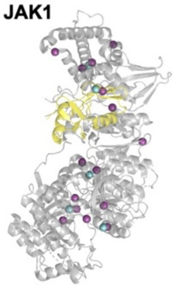

| Residue | Predicted $pK_a$ |
|---|---|
| His509 | 7.1 ± 0.4 |
| His183 | 7.3 ± 0.0 |
| Glu924 | 7.3 ± 0.0 |
| Glu666 | 7.7 ± 0.0 |
| Asp774 | 7.7 ± 0.0 |

D

**JAK2**

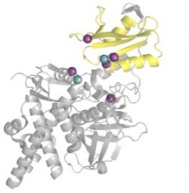

| Residue | Predicted $pK_a$ |
|---|---|
| Glu65 | 6.9 ± 0.2 |
| Cys427 | 7.8 ± 0.1 |

E

**STAT6**

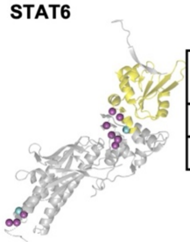

| Residue | Predicted $pK_a$ |
|---|---|
| His145 | 7.3 ± 0.1 |
| Asp519 | 6.8 ± 1.2 |

F

**STAT5A**

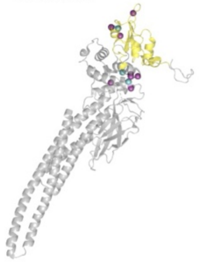

| Residue | Predicted $pK_a$ |
|---|---|
| His476 | 7.1 ± 1.6 |
| His604 | 7.0 ± 0.2 |
| Asp575 | 7.2 ± 2.0 |

G

**CHIO**

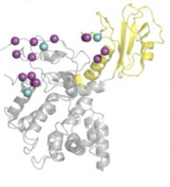

| Residue | Predicted $pK_a$ |
|---|---|
| Glu119 | 7.1 ± 0.0 |
| Cys243 | 7.3 ± 0.0 |
| Asp334 | 7.9 ± 0.0 |

H

**ABL1**

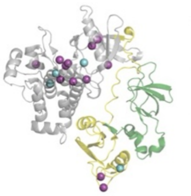

| Residue | Predicted $pK_a$ |
|---|---|
| His220 | 6.9 ± 0.1 |
| Glu450 | 6.6 ± 0.2 |
| Asp400 | 7.9 ±1.7 |
| Asp440 | 6.2 ± 0.4 |

I

**ARHG4**

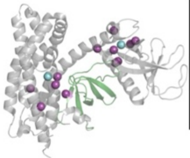

| Residue | Predicted $pK_a$ |
|---|---|
| His526 | 7.2 ± 0.2 |
| Lys426 | 7.0 ± 0.1 |

J

**ACK1**

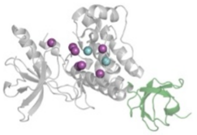

| Residue | Predicted $pK_a$ |
|---|---|
| Asp312 | 6.7 ± 0.0 |
| Glu322 | 7.3 ±0.0 |

K

**AspP2**

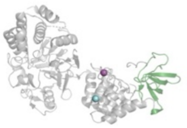

| Residue | Predicted $pK_a$ |
|---|---|
| Cys968 | 6.9 ± 0.1 |

**Cyan- pKa-Shifted Residues**
**Magenta- Ionizing Interactors**
**Yellow- SH2 Domain**
**Green- SH3 Domain**

**Fig. S7. Proteins with modular SH2 domains are predicted to have pH-sensing nodes at the**

**SH2 binding interface, but not at the SH3 domain interface.** Structures of proteins showing the SH2 domain in yellow and the SH3 domain in green. Residues identified through the in silico ionizable network prediction pipeline shown in spheres. Residues with predicted $pK_a$ shifts (cyan) cluster with ionizable interactors (magenta) across the kinase-SH2 domain interaction interface of SHP1 (A), SHP1 (B), ZAP70 (C), JAK1 (D), JAK2 (E), STAT6 (F), STAT5A (G), CHIO (H), ABL1, (I) ARHG4, (J) ACK1, and (K) AspP2.

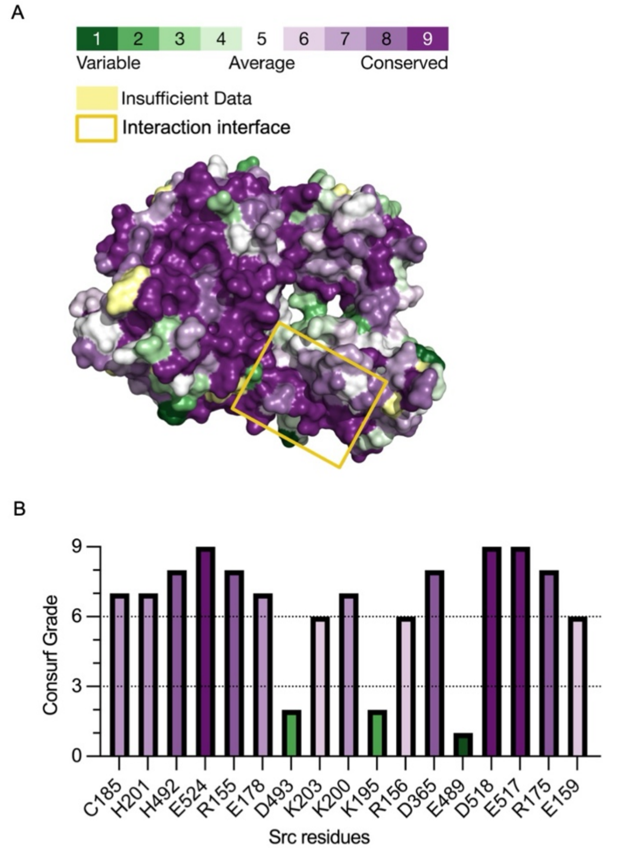


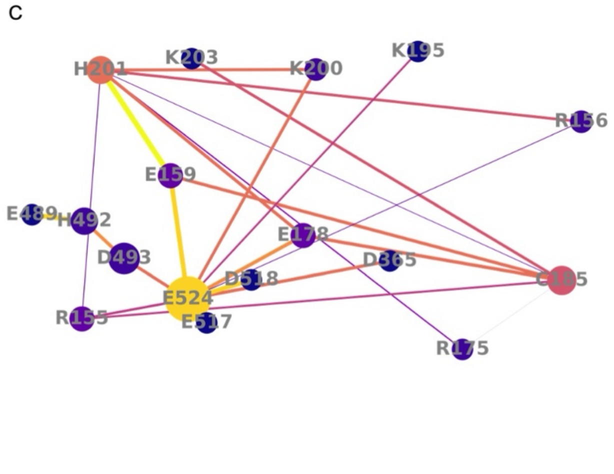


D

**Somatic Src mutations in cancer**

| Residue position | Cosmic Mutation Reference | Amino Acid Mutation | Frequency |
|---|---|---|---|
| C185 | COSM4980479 | C185= | 4/916 |
| H201 | COSM8795277 | H201L | 1/916 |
| H492 | COSM4467372<br>COSM8662426 | H492Q<br>H492= | 2/916 |
| E524 | COSM7430535<br>COSM6506611<br>COSM3389722 | E524K<br>E524D<br>E524K | 9/916 |

=/Coding silent

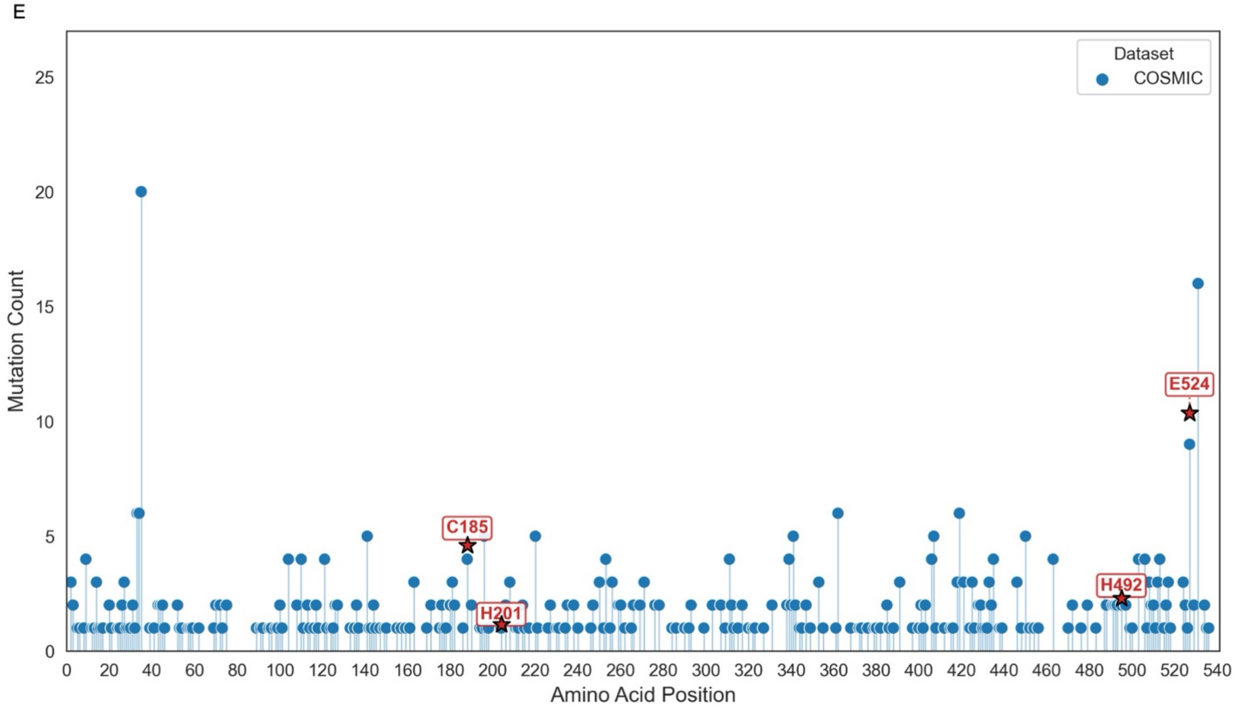


**Fig. S8. Identified residue network in Src is evolutionarily and structurally conserved.** (A)

Consurf surface structure of Src (PDB: 2SRC) with conservation scores for each amino acid from variable to conserved. (B) Consurf grades for shifted residues and their coulombic interactors found in the predicted network. (C) Residue interaction network of shifted ionizable residues and their coulombic interactors. The color spectrum ranges from dark purple to bright yellow. Nodes become more yellow as their degree increases. Larger nodes indicate higher betweenness centrality (measure of how often a node lies on the shortest path between all pairs of nodes in a network). Edges that are yellower and wider represent a higher interaction score. (D) Predicted pH sensing residues and their cosmic mutations. (E) Positions and frequencies of cancer hotspot somatic mutations in Src derived from Catalog of Somatic Mutations in Cancer (COSMIC) database with predicted pKa shifted residues annotated on plot.

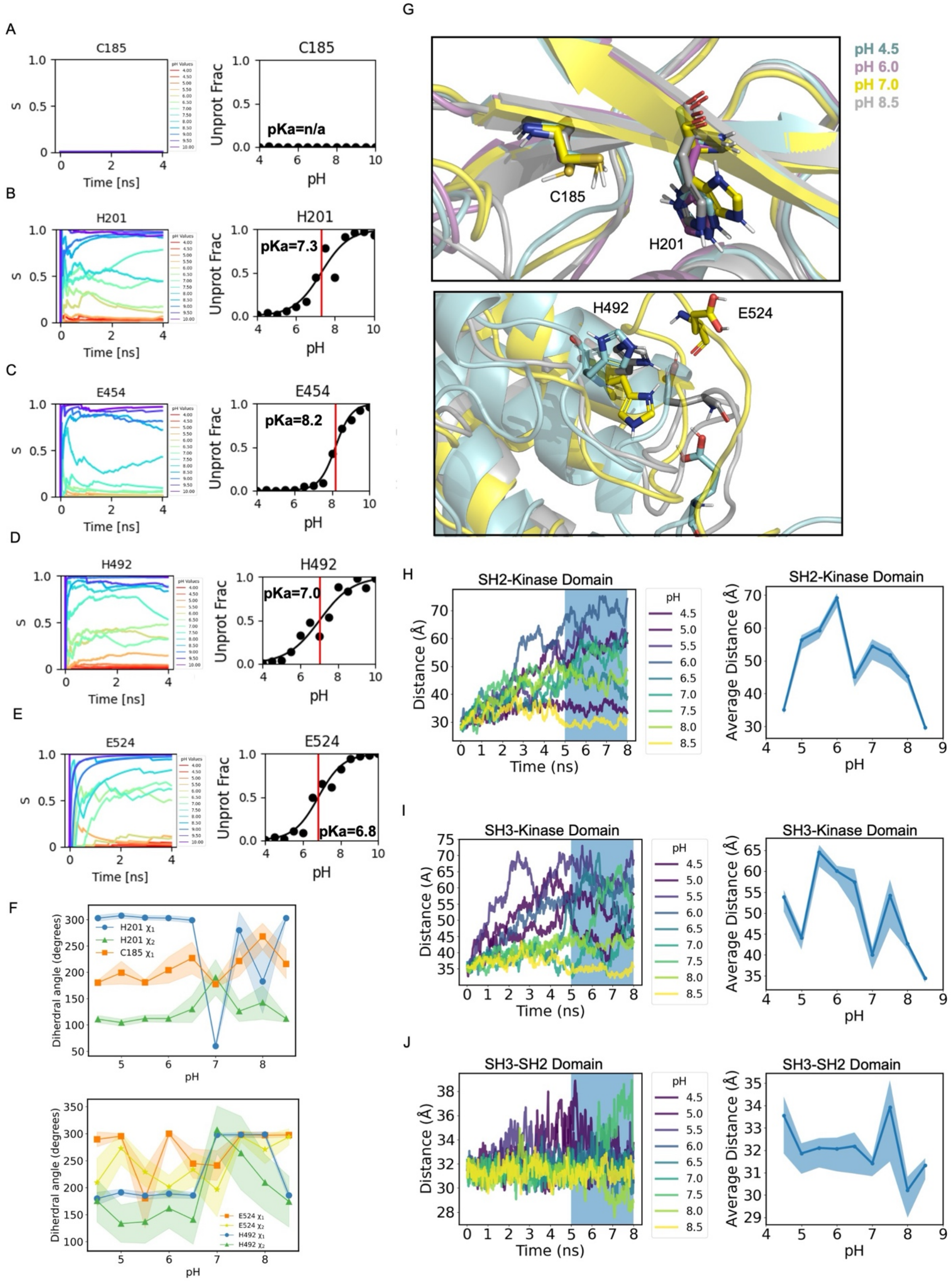
A
C185
S
Time [ns]
pH Values
Unprot Frac
pKa=n/a
pH
B
H201
pKa=7.3
C
E454
pKa=8.2
D
H492
pKa=7.0
E
E524
pKa=6.8
F
Diherdral angle (degrees)
H201 χ1
H201 χ2
C185 χ1
E524 χ1
E524 χ2
H492 χ1
H492 χ2
G
pH 4.5
pH 6.0
pH 7.0
pH 8.5
C185
H201
H492
E524
H
SH2-Kinase Domain
Distance (Å)
Time (ns)
pH
Average Distance (Å)
I
SH3-Kinase Domain
Distance (A)
J
SH3-SH2 Domain

**Fig. S9. Constant pH molecular dynamics (CpHMD) reveals the pH-dependent release of SH2 domain of Src.** (A-E) Time series from CpHMD showing protonation states of indicated Src residues for the first 4ns for pH values 4.0-10.0 (left) and titration plot and $pK_a$ (right) for C185 (A) and H201 (B), E454 (C), H492 (D), and E524 (E). (F) Plot of dihedral angles of C185, H201, H492, and E542 for the last 3ns for pH values 4.5- 8.5. (G) Pair-fit views of zoomed in structures of Src showing the stick views of C185, H201, and H492 at 8ns for pH 4.5 (cyan), 6.0 (magenta), 7.0 (yellow), and 8.5 (grey). (H-J) Time series from CpHMD showing interdomain distances for pH values 4.5-8.5 (left) and average interdomain distance (over the last 3 ns of simulation time) (mean ± SD) (right) for SH2-kinase domains (H), SH3-kinase domains (I), and SH2-SH3 domains (J).

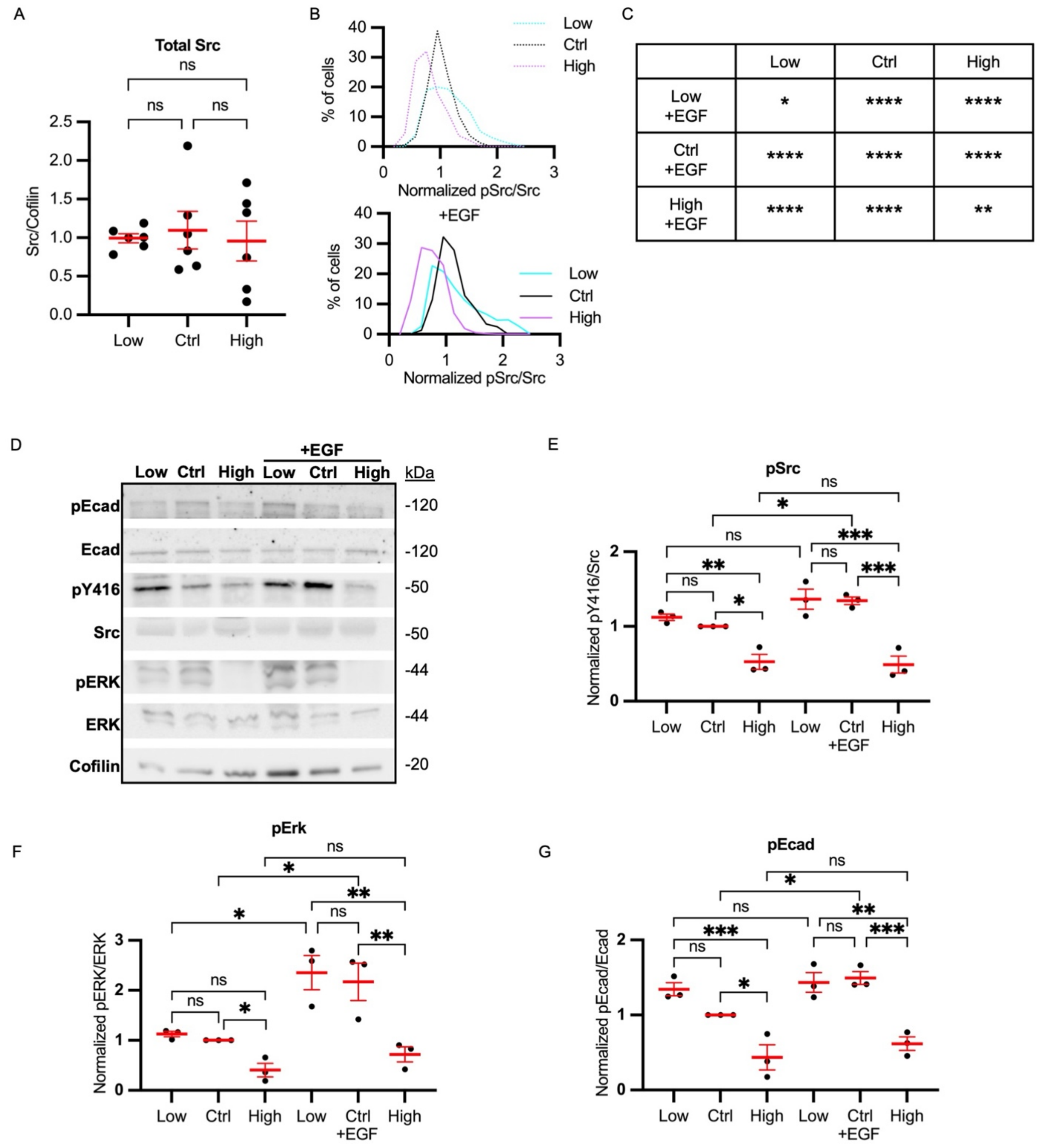


**Fig. S10. pH dependent activation of Src led to downstream signaling activation.** (A) Quantification of Src/Cofilin as described in Fig 5A. (mean ± SEM, N=5). (B) Histogram of single cell data for data shown in Fig. 5F. Low treatment conditions shown in cyan, control in black and high treatment shown in magenta. -EGF above and +EGF below. (C) Table showing Kolmogorov-Smirnov tests comparing distributions of -EGF conditions vs +EGF conditions (D) Representative immunoblot for pSrc (pY416), pERK (pT202/ pY204), pEcad (pY685) after 1 hour of pH treatment and 5 mins of +/- EGF stimulation. (E) Quantification of pY416 src from replicate data collected as in D. Data were normalized to the control in each biological replicate. Scatter plots show (mean ± SEM, N=3). (F) Quantification of pT202/ pY204 ERK from replicate data collected as in D. Data were normalized to the control in each biological replicate. Scatter plots show (mean ± SEM, N=3). (G) Quantification of pY685 Ecad from replicate data collected as in D. Data were normalized to the control in each biological replicate. Scatter plots show (mean ± SEM, N=3). For A, significance was determined using a one-way ANOVA. For C, significance was determined using Kolmogorov-Smirnov tests. For E, F, G significance was determined using one-sample t-test for comparisons to control and one-way ANOVA. * $p<0.05$, **$p<0.01$, ***$p<0.001$, ****$p<0.0001$.

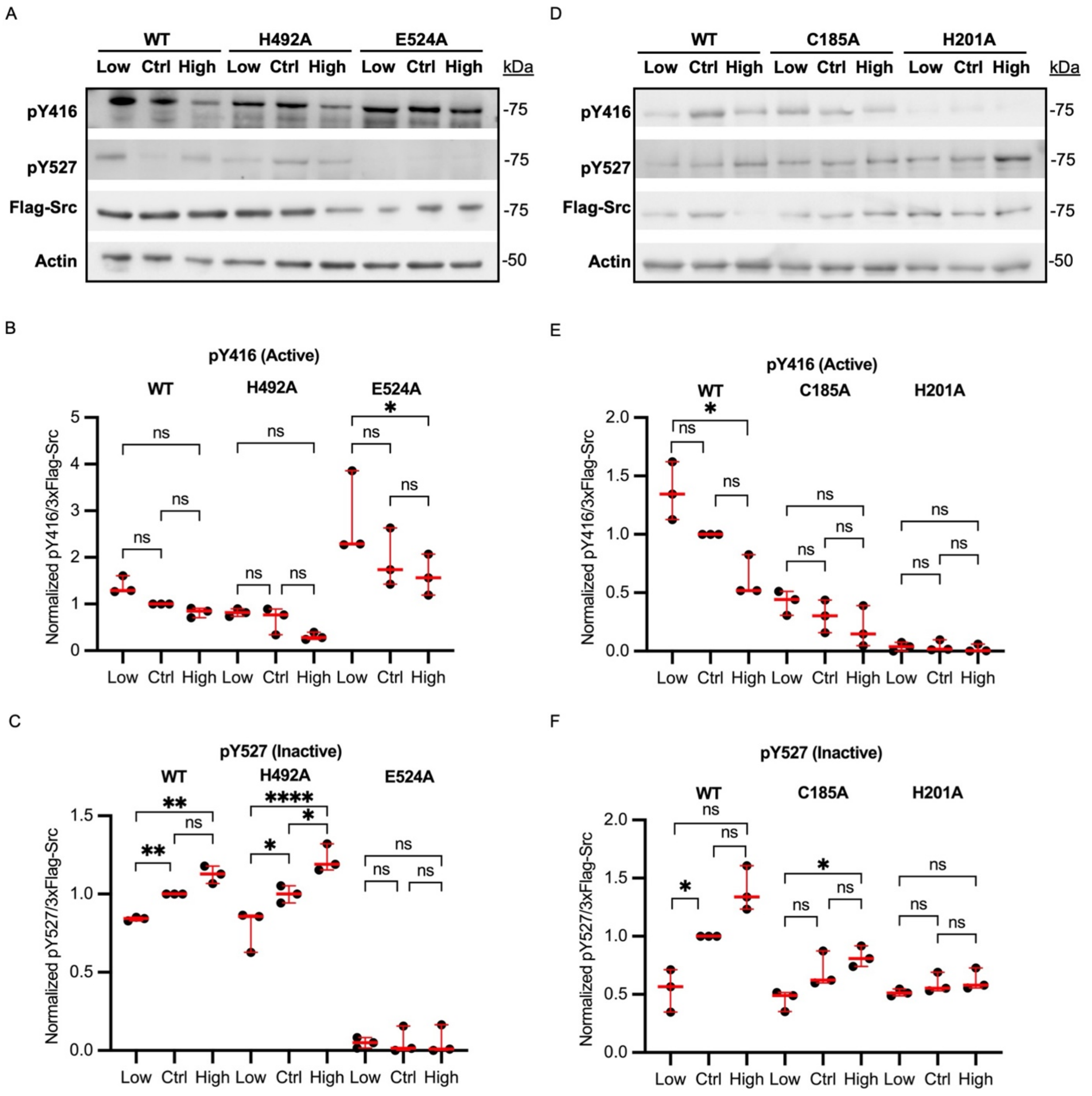


**Fig. S11. Single point mutants reveal distinct contributions to pH-dependent wild-type Src activity.** (A) Representative Western blots of MCF10A lysates overexpressing WT, H492A, and E524A Src. Lysates were prepared from cells maintained for 1 hour at low or high pHi or untreated (control). Immunoblots show total Src and indicated Src phosphorylation sites that reflect protein activation pY527 (inhibited) and pY416 (active). (B-C) Quantification of pY416 (active) (B) and pY527 (inactive) Src (C) from replicate data collected as in A. Data were normalized to WT control in each biological replicate. Scatter plots show (median ± IQR, N=3). (D) Representative immunoblot from MCF10A lysates overexpressing WT, C185A, and H210A Src. Lysates were prepared from cells maintained for 1 hour at low or high pHi or untreated (control). (E-F) Quantification of pY416 (active) (E) and pY527 (inactive) Src (F) from replicate data collected as in D. Data was normalized to WT control in each biological replicate. Scatter plots show (median ± IQR, N=3). For B, C, E, and F, significance was determined using a ratio-paired and one-sample t-test for WT control conditions. For the other mutants, significance was determined using one-way ANOVA. * $p<0.05$, **$p<0.01$, ***$p<0.001$, ****$p<0.0001$”.

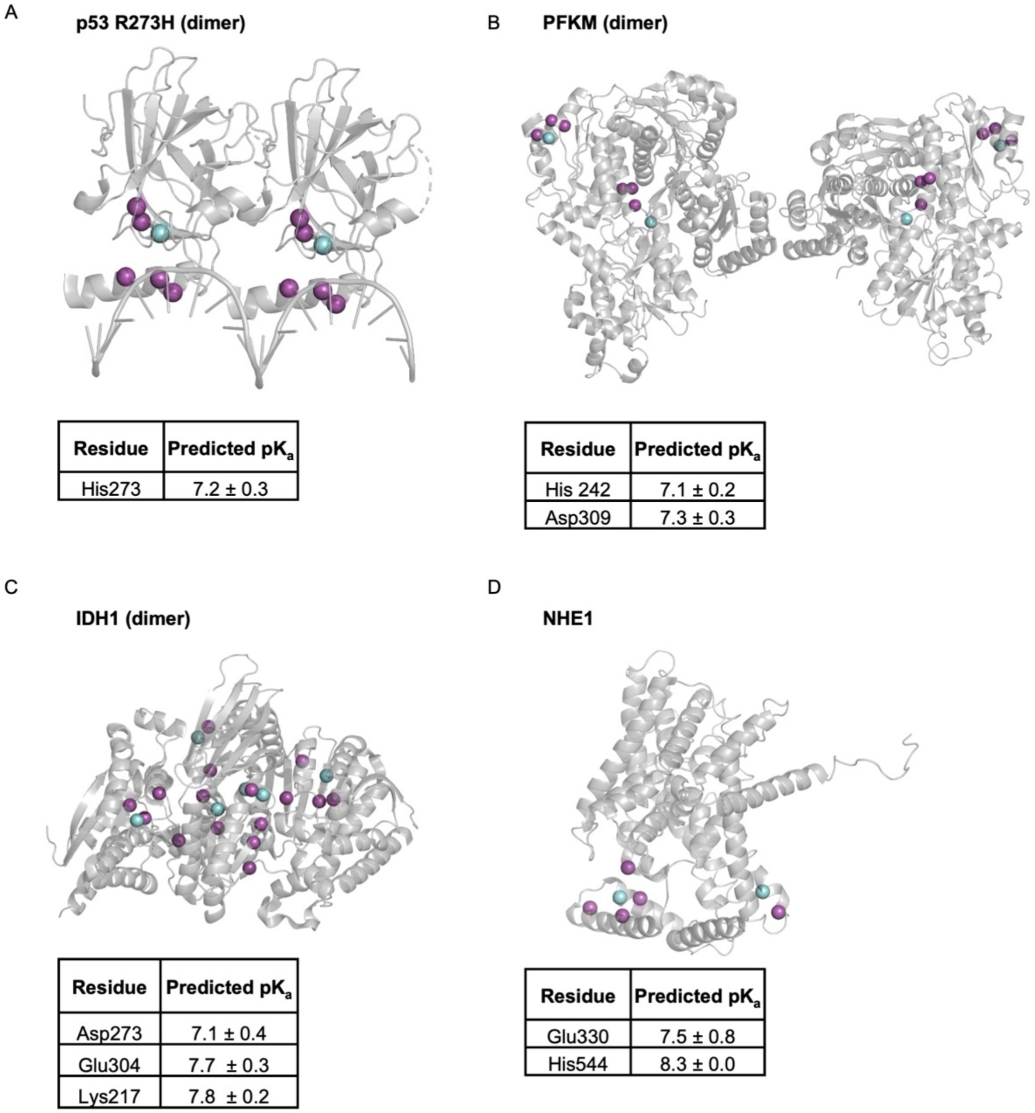


| Residue | Predicted $pK_a$ |
|---|---|
| His273 | 7.2 ± 0.3 |

| Residue | Predicted $pK_a$ |
|---|---|
| His 242 | 7.1 ± 0.2 |
| Asp309 | 7.3 ± 0.3 |

| Residue | Predicted $pK_a$ |
|---|---|
| Asp273 | 7.1 ± 0.4 |
| Glu304 | 7.7 ± 0.3 |
| Lys217 | 7.8 ± 0.2 |

| Residue | Predicted $pK_a$ |
|---|---|
| Glu330 | 7.5 ± 0.8 |
| His544 | 8.3 ± 0.0 |

**Fig. S12. Prediction pipeline accurately identifies pH-sensing nodes of previously characterized pH sensors.** (A) Structure of p53 R273H (PDB: 7B4A) showing residues with predicted $pK_a$ shifts in cyan and coulombic interactors in magenta. (B) Structure of PFKM (PDB: 3O8N) showing residues with predicted $pK_a$ shifts in cyan and coulombic interactors in magenta. (C) Structure of IDH1 (PDB: 1T0L) showing residues with predicted $pK_a$ shifts in cyan and coulombic interactors in magenta. (D) Structure of NHE1 (PDB: 7DSX) showing residues with predicted $pK_a$ shifts in cyan and coulombic interactors in magenta.

**Supplementary Movies Information:**
***Supplementary Movies 1-13: CpHMD Trajectories for SHP2.*** Trajectories are from pH 4.0-10.0 in 0.5 unit increments. Shown in yellow is the N-SH2 domain, orange is C-SH2 domain, and grey is phosphatase domain.

***Supplementary Movies 14-26: CpHMD Trajectories for Src.*** Trajectories are from pH 4.0-10.0 in 0.5 unit increments. Shown in green is SH3 domain, yellow is SH2 domain, and grey is kinase domain.
